# Septin crosstalk with microtubules and actin is regulated by a GSK3-dependent phosphoswitch

**DOI:** 10.64898/2026.05.06.723191

**Authors:** Md Noor A. Alam, TrishaJean C. Holt, Andrew W. Schaefer, Franklin Mayca-Pozo, Sarath Reghunathan, Sarah M. Butts, Priyanka Bhakt, Ilona A. Kesisova, Elias T. Spiliotis

## Abstract

Septins are cytoskeletal filaments that associate with the actin and microtubule cytoskeleton, but the mechanisms that govern septin crosstalk and function with these networks are largely unknown. Here, we show that glycogen synthase kinase 3 (GSK3) directly phosphorylates septin-9 (SEPT9), acting as a molecular switch that bidirectionally controls septin distribution between actin and microtubules. We show that GSK3 inhibition redistributes endogenous SEPT9 toward microtubules in multiple cell types. Phosphomimetic mutations at serines 82 and 85 reduce microtubule binding and enhance actin association in cells and in vitro, while phosphonull mutations promote microtubule binding and growth. In primary hippocampal neurons, GSK3β inactivation promotes SEPT9-microtubule association, and phosphomimetic mutations impair asymmetric neurite growth during neuronal polarization. These findings reveal a phosphorylation-dependent mechanism of septin partitioning between actin and microtubules, placing the cytoskeletal functions of septins under the control of GSK3 – a kinase linked to multiple signaling pathways of cell physiology and metabolism.

**Highlights:**

- GSK3β phosphorylates SEPT9, and its activity gates septin-cytoskeleton association
- S82/S85 phosphorylation reduce microtubule binding and increase actin localization
- Unphosphorylated SEPT9 binds preferentially to microtubules, promoting their growth
- GSK3β inactivation drives SEPT9 to microtubules to establish neuronal polarity

## Introduction

Septins constitute a distinct cytoskeletal network of filaments that interact with both microtubules and actin microfilaments, regulating their organization, dynamics, and crosstalk, and function in cytoskeleton-dependent processes such as membrane traffic, cell division and cell migration ^1–4^. In mammalian cells, septins associate simultaneously with subsets of microtubules and actin filaments ^5,6^. However, the extent of association with each network varies depending on cell type, cellular state, and biological context ^2,5,6^. While septins appear to be homeostatically distributed across cytoskeletal networks, the mechanisms governing their partitioning and redistribution between microtubule and actin systems remain poorly understood. This presents a fundamental knowledge gap in our understanding of how septin organization and function is homeostatically regulated, and how it is coupled to the signaling pathways of cell morphogenesis and physiology.

Septins are GTP-binding proteins encoded by 13 paralogous genes in mammals ^1,7^. They multimerize into non-polar oligomers and polymers comprising subunits from four distinct subgroups (SEPT2, SEPT6, SEPT7, SEPT3) ^7–9^. The fundamental protomeric unit is an octameric complex of two heterotetramers – each consisting of SEPT2, SEPT6, SEPT7 and SEPT9 - that interact a head-to-head manner via SEPT9 ^10–13^. As the central subunit mediating hetero-tetramer dimerization ^12,13^, SEPT9 is the only member of the SEPT3 subgroup that is both ubiquitously expressed and essential for mammalian development ^14–17^; *SEPT9*-null mice die early in gestation ^16^. Notably, SEPT9 mediates septin interactions with both microtubules and actin filaments through a positively charged, intrinsically disordered N-terminal domain ^18–23^. Mutations within this domain underlie the pathogenesis of hereditary neuralgic amyotrophy (HNA), a rare neuropathy ^24^. This ∼150 amino acid region contains microtubule-binding motifs that interact directly with the C-terminal tails of tubulin ^19,22^. Although the same region bundles actin filaments ^20^, it lacks canonical actin-binding motifs and the biochemical basis of this interaction remains unclear. Furthermore, whether and how SEPT9 association with microtubules and actin is regulated has not been established.

Redistribution of septins from actin filaments to microtubules occurs across diverse cell systems and biological processes, and in response to metabolic and pathogenic signals. In the budding yeast *S. cerevisiae*, septins associate with microtubules under nutrient-limiting conditions ^25^. In mammalian cells, the microtubule-stabilizing drug taxol enhances septin-microtubule binding by suppressing expression of Cdc42 effector proteins 2/3 (Cdc42EP2/3), which otherwise facilitate septin-actin binding ^26^. Additionally, actin depolymerization induced by *Chostridium difficile* transferase promotes septin association with microtubule plus ends ^27^, which is also enhanced in cells infected with the hepatitis C virus ^28^. Conversely, septins translocate to stress fibers following stimulation with lysophosphatidic acid - an activator of G protein-coupled receptors - and after microtubule depolymerization ^22,29,30^. The apparent competition between microtubules and actin filaments for a limiting septin pool is consistent with the observation that intracellular tubulin and actin concentrations are 10- to 100-fold higher than those of septins ^31–33^.

Despite the functional importance of septin-cytoskeleton interactions, the signaling pathways and molecular mechanisms that govern septin association with microtubules and actin are ill defined. Septin assembly is regulated through Rho signaling. In budding yeast, septin assembly requires cycles of Cdc42 activation, and depends on the Cdc42 effector proteins (Gic1/2), which crosslink septin filaments ^34–38^. In mammalian cells, Cdc42 activity maintains septin localization to actin stress fibers, and the Cdc42 effector protein 3 (CdcEP3; Borg2) promotes actin filament assembly when in complex with septins ^26,39–41^. RhoA activation induces septin assembly ^42^, and septin associations with the Rho guanine exchange factors ARHGEF18 and the Rho effector Rhotekin suggest a feedback loop between Rho signaling and actin assembly through septins^29,43–45^. Septin-actin interactions are further modulated by mechanical signals transduced in response to extracellular matrix stiffness and porosity, but the molecular mechanisms are unclear ^39,46–48^.

Regulation of septin-microtubule interactions is even less well understood. Microtubule-septin binding is regulated indirectly by microtubule-associated proteins (MAPs; e.g., MAP4) which compete with septins for microtubule association, and/or by Cdc42 inactivation and Cdc42EP3 downregulation – both which reduce septin-actin affinity ^26,49–51^. However, mechanisms of direct regulation have yet to be identified. Phosphorylation of MAPs by signaling kinases is a well-established paradigm for controlling microtubule-binding ^52^. Given that SEPT9-microtubule interactions are electrostatic in nature - involving basic residues in SEPT9 N-terminus and the acidic and polyglutamylated C-terminal tails of polymerized tubulin ^19,21,51,53^, we hypothesized that SEPT9 phosphorylation may directly regulate septin-microtubule binding.

Although phosphorylation of various septin paralogs has been shown to regulate their assembly, subcellular localization, and interaction with binding partners ^54,55^, phospho-regulation of septin interactions with the actin or microtubule cytoskeleton has not previously been demonstrated. Here, we investigate whether phosphorylation of two N-terminal serine (Ser; S) residues of SEPT9 – S82 and S85 – regulate septin association with microtubules and actin filaments. Previous analysis of the phosphoproteome of the glycogen synthase kinase 3 (GSK3) – a ubiquitously expressed kinase that regulates the microtubule-binding of several MAPs – uncovered S85 and S82 of SEPT9 as putative phosphorylation residues ^56,57^. Here, we show that phosphorylation at these sites acts as a molecular switch, reducing septin affinity for microtubules while enhancing association with actin filaments.

## Results

### SEPT9 is a direct and physiological target of GSK3β

Multiple signaling pathways — including Wnt, ERK, and AKT — converge on the GSK3 signaling cascade to regulate cell morphogenesis, proliferation, metabolism, and cell-fate determination ^56,58,59^. Because GSK3 is constitutively active, inhibition by upstream kinases serves as a permissive "on-switch" for a broad range of downstream molecular events, including the regulation of cytoskeletal organization through small GTPases and cytoskeletal-associated proteins such as tau and CLASP, whose affinity for the cytoskeleton is directly modulated by GSK3-mediated phosphorylation ^56,60–62^.

Both GSK3 isoforms, GSK3α and GSK3β, phosphorylate the first serine or threonine of the pentapeptide motif S/T-X-X-X-S/T, where X denotes any amino acid ^62^. This requires prior phosphorylation of the S/T residue at the +4 position — a prerequisite known as priming phosphorylation ^62^. In an unbiased quantitative phosphoproteomic screen for GSK3 substrates using embryonic stem cells (ESCs) from wild-type and *Gsk3* double-knockout mice, S82 and S85 of SEPT9 were identified as putative phosphosites ^57^.

S82 and S85 are located in the N-terminal region of SEPT9 that precedes the GTP-binding domain (Figure 1A). This region is enriched in positively charged residues and has accordingly been termed the "basic domain" ^19^. Notably, it interacts directly with microtubules and actin filaments ^19,20^. Both phosphosites are flanked by two clusters of K/R-X-X-D/E motifs, which collectively constitute a minimal sequence – aa 61-113 of SEPT9 isoform 1 (SEPT9_i1) – that can directly bind and bundle microtubules^19^. S82 and S85 are present in the three longest SEPT9 isoforms (isoforms 1, 2, and 3), which differ primarily in the length of their N-terminal extension (the first ∼25–37 residues), and are conserved across human, mouse, rat and canine SEPT9 sequences despite considerable divergence in the N-terminal region (Figure 1B). Both residues are absent from the shorter isoforms 4 and 5, which lack most of the N-terminal extension (Figure 1A).

**Figure 1.**
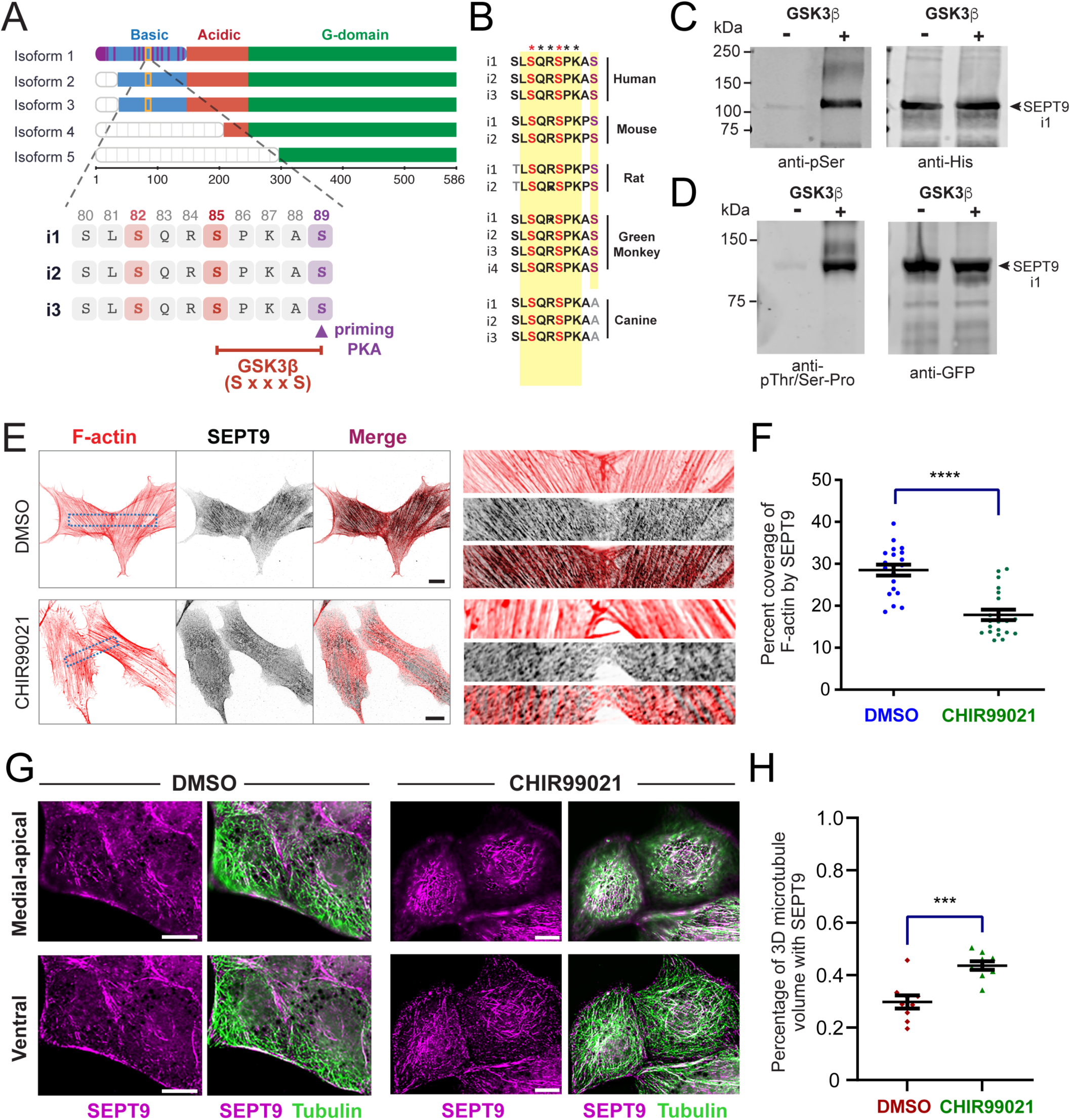
GSK3 directly phosphorylates SEPT9 and regulates SEPT9 localization to actin stress fibers and microtubules. **(A)** Schematic of the domains of the five human SEPT9 isoforms, and alignment of the amino acid sequences containing the putative GSK3 phosphorylation sites in the three longest isoforms (i1, i2, i3). Serine residues (S82, S85) identified as GSK3 phosphorylation targets in SEPT9_i1 are highlighted in pink. Microtubule-binding motifs are shown in dark purple in the longest SEPT9_i1 isoform. The priming serine residue is highlighted in light purple. **(B)** Multiple sequence alignment of SEPT9 isoforms from human, mouse, rat, green monkey and canine organisms, illustrating conservation of the putative GSK3 phosphoserine residues (red). Conserved residues are highlighted in yellow. Asterisks denote fully conserved positions. **(C)** Western blots of recombinant His-GFP-SEPT9_i1 following an *in vitro* kinase reaction in the presence or absence of GSK3β, probed with antibodies against phosphoserine/threonine (left) and His (right). **(D)** Western blots of recombinant His-GFP-SEPT9_i1 following an *in vitro* kinase reaction in the presence or absence of GSK3β, probed with antibodies against the phospho-threonine/serine-proline motif (left) and GFP (right). **(E–F)** Super-resolution SoRa spinning-disk confocal microscopy images (E) show single ventral optical sections of U2OS cells co-stained with anti-SEPT9 (inverted red) and phalloidin (inverted grayscale) following treatment with CHIR99021 (10 μM) or DMSO (control) for 12 hours. Dashed rectangles indicate regions shown at higher magnification. Scale bar, 10 μm. Quantification (F) of the percentage (mean ± SEM) of phalloidin-segmented actin filament area co-occupied by SEPT9 fluorescence signal (*n* = 20 cells). Statistical comparisons were made using a Mann–Whitney U test. ****, p < 0.0001. **(G–H)** Denoised and deconvolved wide-field fluorescence microscopy images (G) show maximum intensity projections of medial-apical (top) and ventral (bottom) optical sections of MDCK cells stained for SEPT9 (magenta) and α-tubulin (green) following treatment with CHIR99021 (10 μM) or DMSO (control) for 2 hours. Scale bar, 10 μm. Quantification (H) of the percentage (mean ± SEM; *n* = 8) of the 3D microtubule volume that overlaps with SEPT9. Statistical comparisons were made using a Mann–Whitney U test. ***, p < 0.001.

S85 conforms to the canonical GSK3 phosphorylation motif S-X-X-X-S, with S89 at the +4 position serving as a putative priming site ^62^. Notably, the +2 position relative to S85 is occupied by a proline residue, which is preferentially recognized by GSK3β. This proline may also confer additional regulatory complexity as the pSer-Pro motif is a known substrate for prolyl isomerases, which can modulate downstream protein interactions and inhibit dephosphorylation through *cis/trans* isomerization ^63,64^. By contrast, S82 does not conform to the canonical GSK3β consensus because the nearest downstream serine (S85) is only three residues away — one position short of the n+4 spacing required to fit the structural docking geometry of the GSK3β priming-phosphate binding pocket ^62^. Its detection in the phosphoproteomic screen therefore may reflect a non-primed phosphorylation event or the activity of a distinct upstream kinase.

To validate SEPT9 as a direct GSK3 substrate, we performed in vitro kinase assays using recombinant SEPT9_i1 purified from bacteria together with recombinant GSK3β, and independently assessed the impact of GSK3 activity on SEPT9 localization in cells. In vitro kinase reactions of GSK3β with SEPT9 resulted in robust phosphorylation, detected with a phosphoserine (pSer) antibody (Figure 1C) and independently corroborated with a phospho-threonine/serine-proline (pThr/Ser-Pro) antibody that recognizes Ser-Pro motifs such as that of S85 (Figure 1D). Consistent with a direct phosphorylation of SEPT9 by GSK3β, pharmacological inhibition of GSK3 with CHIR99021 — a highly selective and potent inhibitor ^65,66^ — altered the subcellular localization of endogenous SEPT9 in two different cell types. In human osteosarcoma U2OS cells, where endogenous SEPT9 localizes predominantly to actin stress fibers ^22,23^, super-resolution SoRa spinning disk confocal microscopy revealed that CHIR99021 reduced SEPT9 colocalization with ventral stress fibers (Figure 1E-F). Conversely, in renal MDCK epithelial cells, where SEPT9 associates primarily with microtubules ^19,51,67^, volumetric colocalization analysis of MDCK image stacks demonstrated increased SEPT9 association with the microtubule network following GSK3β inhibition (Figure 1G-H). Collectively, these findings from unbiased phosphoproteomics ^57^, in vitro kinase assays, and pharmacological inhibition establish SEPT9 as both a direct and physiological GSK3 substrate, and suggest that GSK3-mediated phosphorylation bidirectionally regulates SEPT9 partitioning between the actin and microtubule cytoskeletons.

### Phosphonull and phosphomimetic mutations of SEPT9_i1 at S82/S85 differentially regulate association with actin stress fibers and microtubules

Given that pharmacological inhibition of GSK3β reduced septin colocalization with actin stress fibers in U2OS cells, we sought to determine whether SEPT9_i1 association with actin stress fibers is directly regulated by phosphorylation at S82 and S85, the putative GSK3 phosphorylation sites. We generated single phosphonull (S-to-A) and phosphomimetic (S-to-E) mutations at the S85 residue of GFP-SEPT9_i1, as well as double mutations that combined S82, which along with S85 was identified in the phosphoproteome of GSK3 ^57^. To minimize epistasis from endogenous SEPT9, we used a plasmid co-expressing an shRNA targeting all endogenous SEPT9 isoforms and an shRNA-resistant GFP-SEPT9_i1. As previously validated in HeLa cells ^68^, expression of this plasmid depletes endogenous SEPT9 isoforms — confirmed by immunolabeling with a pan-isoform anti-SEPT9 antibody — without affecting the expression of shRNA-resistant variants. Endogenous SEPT9 was similarly knocked down in U2OS and MDCK cells (Figure S1A-B).

Consistent with previously reports of SEPT9 localization in multiple cell types ^19,23,69–72^, wild-type GFP-SEPT9_i1 appeared as fine, elongated fibers colocalizing with ventral actin stress fibers, and as thicker bundle-like filaments colocalizing with a subset of perinuclear microtubules in the medio-apical cytoplasm (Figure 2A). In 53% of cells, SEPT9_i1 was also detected in the nucleus, which is in agreement with prior observations and may reflect a cell cycle-dependent nuclear import driven by the N-terminal nuclear localization signal of SEPT9 ^69,73^. In maximal projections of medio-apical confocal sections acquired by spinning disk microscopy, the thick microtubule-associated bundles of GFP-SEPT9_i1 were markedly diminished or completely absent by the single (S85E) and double (S82E/S85E) phosphomimetic mutations (Figure 2B and S1C). Instead, the phosphomimetic mutants were highly enriched on ventral actin stress fibers, as visualized in maximal projections from the two most ventral optical sections (Figure 2B and S1C). Quantification of the percentage of ventral stress fiber area occupied by GFP-SEPT9_i1 confirmed a greater than 2-fold increase for both phosphomimetic mutants relative to wild-type (Figure 2C-D). Notably, in cells which expressed the phosphomimetic mutants, the ventral actin stress fiber network spanned a larger fraction of the total surface area, which was consistent with a role of SEPT9_i1 in stabilizing actin stress fibers ^23,74–76^ (Figure 2E-F). Nuclear localization of GFP-SEPT9_i1 S85E and S82E/S85E was comparable to wild type (51% vs. 53%; *n* = 40–42), indicating that phosphorylation at these sites selectively affects cytoskeletal — rather than nuclear — targeting.

**Figure 2.**
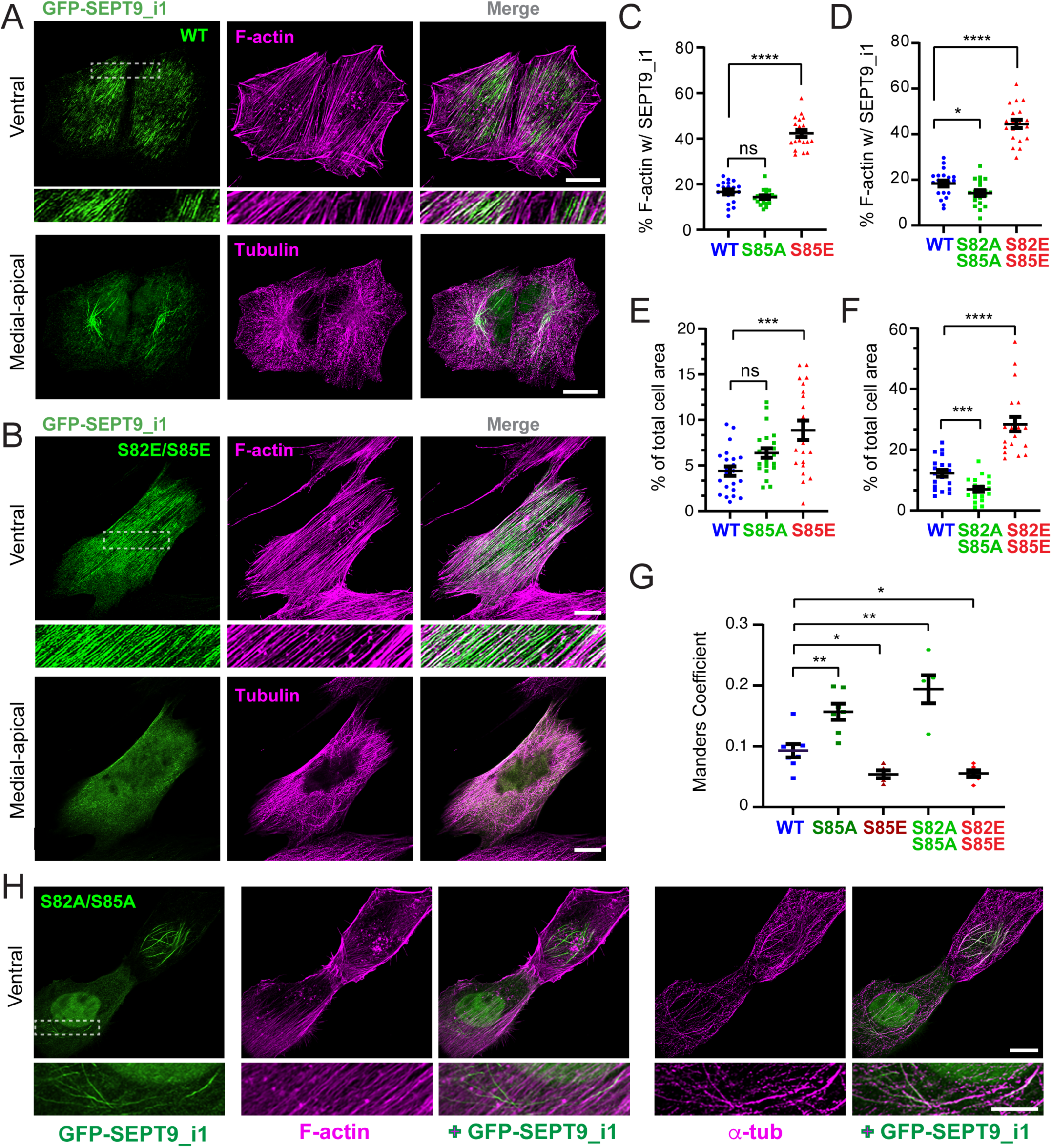
Phosphomimetic and phosphonull mutations of SEPT9 at S82 and S85 bidirectionally alter colocalization with actin stress fibers and microtubules. **(A-B)** Super-resolution SoRa spinning-disk confocal microscopy images show maximum intensity projections of ventral (top) and medial-apical (bottom) optical sections of U2OS cells transfected with wild-type (WT) GFP-SEPT9_i1 (green; A) or GFP-SEPT9_i1-S82E/S85E (green; B), stained for actin (phalloidin; magenta) and α-tubulin (magenta). Dashed rectangles indicate regions shown at higher magnification. Scale bars, 10 μm. **(C-D)** Quantification of the percentage (mean ± SEM) of phalloidin-segmented actin filament area occupied by SEPT9 fluorescence signal in single ventral optical sections of U2OS cells (*n* = 20) transfected with plasmids expressing SEPT9 shRNA and shRNA-resistant wild type (WT) GFP-SEPT9_i1, GFP-SEPT9_i1-S85A or GFP-SEPT9_i1-S85E (C), and GFP-SEPT9_i1-S82A/S85A or GFP-SEPT9_i1-S82E/S85E (D). Pair-wise statistical analyses were made with unpaired Welch’s t-test; Mann-Whitney U-test was made for comparison of WT to S85A mutant in (C). *, p < 0.05; ****, p < 0.0001; ns, not significant **(E-F)** Quantification of phalloidin-segmented actin filament area as a percentage (mean ± SEM) of total surface area in single ventral optical sections of U2OS cells (*n* = 20) transfected with plasmid expressing SEPT9 shRNA and shRNA-resistant wild type (WT) GFP-SEPT9_i1, GFP-SEPT9_i1-S85A or GFP-SEPT9_i1-S85E (E), and GFP-SEPT9_i1-S82A/S85A or GFP-SEPT9_i1-S82E/S85E (F). Pair-wise statistical analyses were made using an unpaired Welch’s t-test; a Mann-Whitney U-test was made for comparison of WT to S82E/S85E mutant in (F). ***, p < 0.001; ****, p < 0.0001; ns, not significant **(G)** Quantification of the Manders’ coefficients (mean ± SEM) of overlap between the GFP-SEPT9_i1 (wild type and phosphomutants) and microtubule fluorescence signals in 3D medial-apical image volumes of U2OS cells imaged by super-resolution SoRa spinning-disk confocal microscopy. Pairwise statistical comparisons were made using a Mann–Whitney U test (*n* = 5-8). *, p < 0.05; **, p < 0.01 **(H)** Super-resolution SoRa spinning-disk confocal microscopy images show maximum intensity projections of ventral optical sections of U2OS cells transfected with GFP-SEPT9_i1-S82A/S85A (green) stained for actin (phalloidin; magenta) and α-tubulin (magenta). Dashed rectangles indicate regions shown at higher magnification. Scale bars, 10 μm.

The intracellular distribution of the phosphonull mutants S85A and S82A/S85A differed from both wild-type SEPT9_i1 and the phosphomimetic variants. Both mutants were characterized by the formation of elongated thick filaments along the nuclear envelope rim, visible in both medio-apical and ventral optical sections — consistent with the substantial thickness of these structures, which colocalized with perinuclear microtubule bundles (Figure 2H and S1D). To avoid non-specific overlap of microtubules with nuclear or actin stress fiber-associated SEPT9_i1, we quantified SEPT9_i1 colocalization with microtubules in the 3D volume regions from the medio-apical cytoplasm of U2OS cells. Colocalization with microtubules increased markedly for both phosphomimetic mutants – a nearly 2-fold increase for the double S82A/S85A mutant (Figure 2G), which in contrast occupied less actin stress fibers and reduced the ventral actin network (Figure 2C-E). Additionally, a higher proportion of cells expressing the phosphonull double mutant S82A/S85A exhibited nuclear localization (64% vs. 53%; n = 40–45), suggesting that in its unphosphorylated state SEPT9_i1 may traffic more efficiently into the nucleus due to either enhanced association with importin-α7 and/or less association with actin stress fibers. Collectively, these data suggest that phosphorylation of S82/S85 by GSK3 promotes selective enrichment on actin stress fibers, while the unphosphorylated state favors microtubule association.

### Phosphorylation of SEPT9_i1 at S82/S85 regulates septin complex recruitment to centrosomal microtubules, and impacts microtubule nucleation and elongation

SEPT9 has been established as a septin paralog that mediates microtubule association with septin complexes, and the SEPT9_i1 isoform in particular not only interacts specifically with microtubules but also regulates microtubule growth ^19,22,77^. We therefore examined whether the phosphomimetic and phosphonull S82/S85 mutations affect septin association with nascent microtubules growing from the centrosome. For these experiments, we used MDCK cells, in which endogenous septins are more prominently associated with microtubules than with actin stress fibers, providing an orthogonal cellular context relative to U2OS cells for analyzing the SEPT9_i1 phosphomutants.

In MDCK cells expressing wild-type or phosphomutant SEPT9_i1 in an endogenous SEPT9 depletion background, microtubules were depolymerized by nocodazole treatment for 2 hours, followed by nocodazole washout and 40 seconds of regrowth. Consistent with endogenous SEPT9 localization to the centrosome and centrosome-proximal segments of the nascent microtubule lattice after nocodazole wash-out in the absence of SEPT9 knock-down (Figure S2A), wild-type GFP-SEPT9_i1 decorated the lattice of centrosomal microtubules (Figure 3A). Both the single (S85E) and double (S82E/S85E) phosphomimetic mutants showed a dramatic reduction in centrosomal microtubule coating, with GFP signal largely absent from the microtubule lattice (Figure 3A-C). Among the phosphonull mutants, the single S85A mutant showed no significant change, while the double S82A/S85A mutant exhibited a statistically significant increase in the fraction of centrosomal microtubules decorated with SEPT9_i1 (Figure 3A-C). Notably, endogenous SEPT2 and SEPT7 mirrored these changes (Figure S3). Both paralogs showed increased association with centrosomal microtubules in cells expressing the double phosphonull mutant (Figure S3A, S3C, and S3E-F), and were markedly depleted from regrown microtubules in cells expressing the double phosphomimetic mutant (Figure S3A, S3D and S3E-F). This indicates that the phosphomutants do not function as monomers or homomeric assemblies, but rather engage microtubules as part of canonical heteromeric septin complexes, and SEPT9 association with microtubules impacts the localization of its heteromeric septin partners.

**Figure 3.**
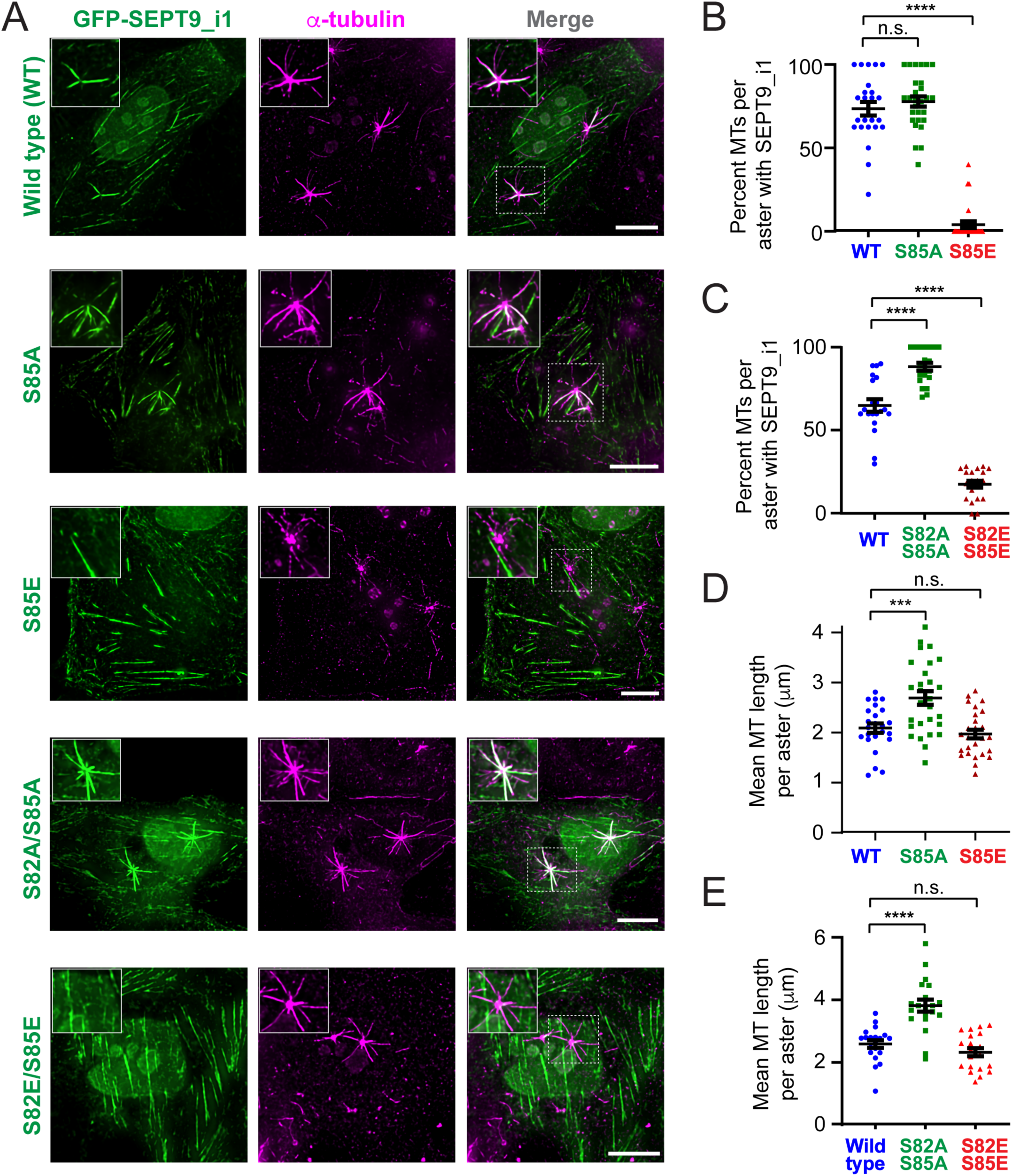
Phosphonull mutations of SEPT9_i1 at S82 and S85 enhance centrosomal microtubule association and growth, whereas phosphomimetic mutants impair microtubule binding. **(A)** Denoised and deconvolved wide-field fluorescence microscopy images show maximum intensity projections of medial-apical optical sections of MDCK cells, which were transfected with plasmids expressing SEPT9 shRNA and shRNA-resistant GFP-SEPT9_i1 (wild-type and phosphomutants; green), and stained for α-tubulin (magenta). Cells were treated for nocodazole (1.6 μM) for 2 h at 37 °C followed by washout for 40 seconds at 25 °C to allow centrosomal microtubule regrowth prior to fixation and staining. Insets show magnified views of regions outlined by dashed rectangles. Scale bars, 10 μm. **(B)** Quantification of the percentage (mean ± SEM) of astral microtubules decorated with GFP-SEPT9_i1 per aster following nocodazole wash-out. Pairwise comparisons between wild-type (*n* = 24 cells), S85A (*n* = 28 cells), and S85E (*n* = 27 cells) mutant were statistically analyzed using a Mann-Whitney U-test. ****, p < 0.0001; n.s., not significant. **(C)** Quantification of the percentage (mean ± SEM) of astral microtubules decorated with GFP-SEPT9_i1 per aster (*n* = 20 cells) after nocodazole wash-out. Pairwise comparisons between the wild-type, S82A/S85A, and S82E/S85E were statistically analyzed with a Mann-Whitney U-test. ****, p < 0.0001 **(D)** Quantification of mean (mean ± SEM) astral microtubule length per aster following nocodazole wash-out in MDCK cells expressing GFP-SEPT9_i1. Pairwise comparisons between the wild-type (*n* = 24 cells), S85A (*n* = 28 cells), and S85E (*n* = 27 cells) mutant were statistically analyzed with Welch’s t-test. ***, p < 0.001; n.s., not significant. **(E)** Quantification of astral microtubule length (mean ± SEM) per aster following nocodazole wash-out in MDCK cells (*n* = 20 cells) expressing GFP-SEPT9_i1. Pairwise comparisons between the wild-type, S82A/S85A, and S82E/S85E were statistically analyzed with an unpaired Welch’s t-test. ****, p < 0.0001; n.s., not significant.

We next analyzed the lengths and numbers of microtubules in centrosomal asters to assess the functional impact of the phosphomutants on microtubule growth. Both single and double phosphonull mutants increased the mean microtubule length per aster and the number of microtubules per aster, indicative of enhanced nucleation and elongation from the centrosome (Figure 3D-E and S2B-C). In contrast, neither the single nor double phosphomimetic mutant affected mean microtubule length or aster microtubule number (Figure D-E, and S2B-C). Thus, the unphosphorylated form appears to promote septin-microtubule association and centrosomal microtubule growth, while the phosphorylated form is excluded from microtubules, redirecting septin complexes away from the microtubule network and is dispensable for microtubule growth.

### Phosphomimetic mutations S82E and S85E reduce in vitro microtubule binding of SEPT9_i1 and SEPT2/6/7/9_i1 complexes

Because disruption of septin colocalization with microtubules and actin by phosphomutations may arise from indirect effects, we directly tested the impact of these mutations using in vitro total internal reflection fluorescence (TIRF) microscopy binding assays. We generated phosphonull (S82A/S85A) and phophomimetic (S82E/S85E) mutations in recombinant GFP-SEPT9_i1, as well as unlabeled SEPT9_i1 in complex with msGFP-SEPT2, SEPT6 and SEPT7, which were expressed in and purified from bacteria ^78^. Prior to assessing microtubule binding, we examined whether these mutations affected the formation and higher-order assembly of the octameric complex (Figure S4). Size exclusion chromatography of affinity-purified msGFP-SEPT2/6/7/9_i1 complexes showed that wild type, S82A/S85A and S82E/S85E preparations each eluted as a single peak in the same volume fractions, corresponding to the predicted size of the hetero-octamer (Figure S4A-C). Western blot analysis of the pooled fractions confirmed the presence of all four subunits in each preparation (Figure S4A-C). Sedimentation assays performed after incubation in low salt buffer, which induces septin assembly into higher order multimers^79^, showed that an equivalent percentage of all three complexes forming higher order polymers (Figure S4D-E). These results indicate that the S82/S85 phoshomutants do not impair SEPT9_i1 folding, hetero-octamer formation, or the assembly-competence of SEPT2/6/7/9_i1 complexes.

We next performed in vitro decoration assays using taxol-stabilized microtubules immobilized on glass chambers. Consistent with previous findings ^77,80^, wild-type SEPT9_i1 bound microtubules at nanomolar concentrations. Quantification of the microtubule-bound GFP-SEPT9_i1 at 1 nM and 3 nM showed a marked reduction in the binding for the phosphomimetic mutant S82E/S85E mutant compared to wild type at both concentrations (Figure 4A-B and S5A-B). At 1 nM, the phosphonull mutant S82A/S85A showed a modest increase in binding relative to wild type, and at 3 nM this difference became more pronounced, with mean microtubule-bound intensity increasing nearly two-fold, while microtubule binding of the phosphomimetic SEPT9_i1_S82E/S85E was as minimal as to the 1 nM condition (Figure 4A-B and S5A-B). The S82E/S85E mutant also substantially reduced microtubule association of the msGFP-SEPT2/6/7/9_i1 complex, whereas SEPT9_i1_S82A/S85A resulted in a statistically significant increase (Figure 4C-D). Consistent with the reduced colocalization of phosphomimetic mutants of SEPT9_i1 with cellular microtubules (Figure 2G and 3A-C) and the enhanced microtubule colocalization of SEPT9 following pharmacological GSK3β inhibition (Figure 1G-H), these in vitro data indicate that phosphorylation of SEPT9_i1 at S82/S85 reduces the microtubule-binding capacity of SEPT9-containing septin complexes.

**Figure 4.**
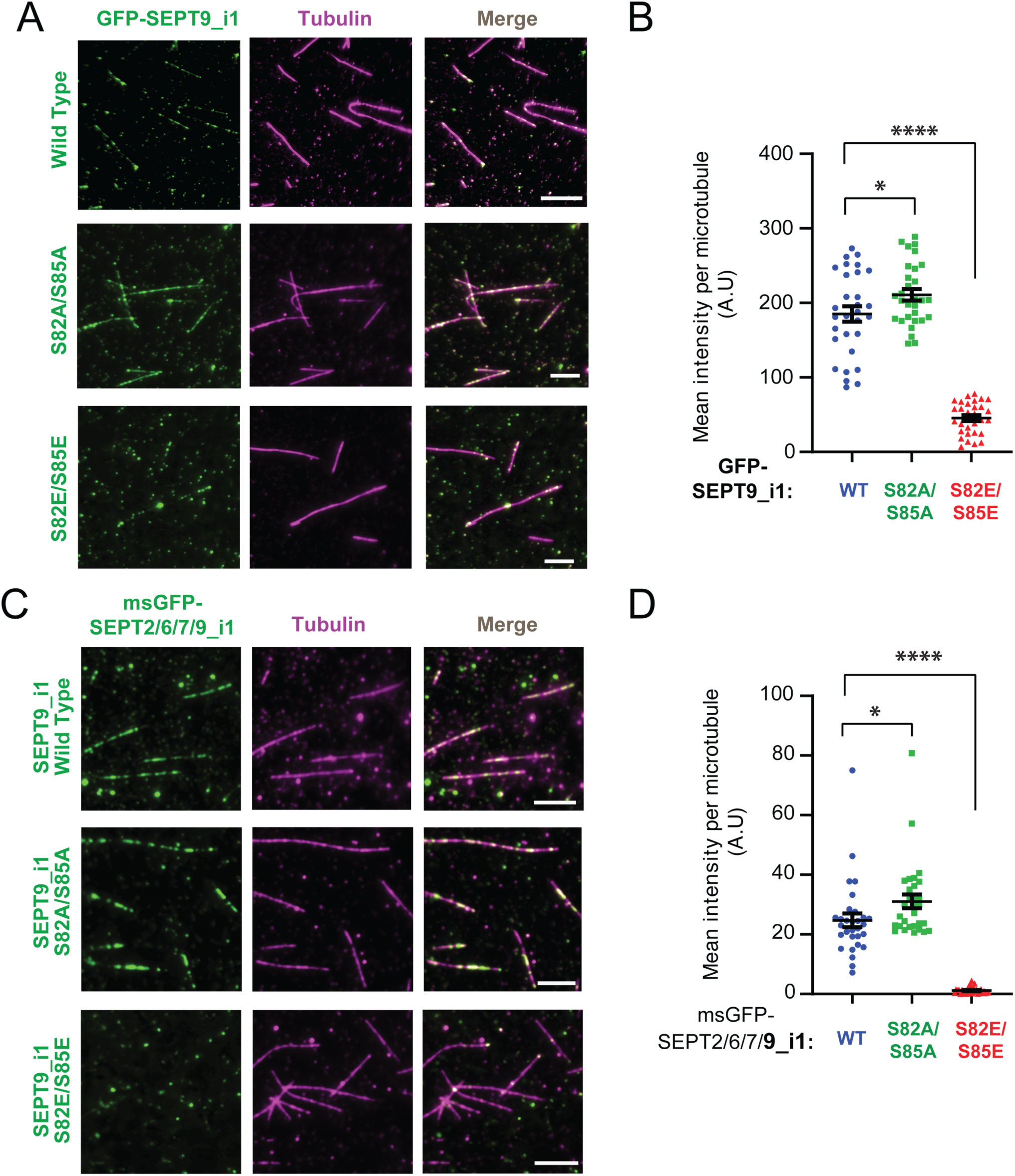
Phosphonull and phosphomimetic mutations at S82/S85 of SEPT9_i1 enhance and abrogate microtubule binding in vitro, respectively. **(A)** Total internal reflection fluorescence (TIRF) microscopy images of taxol-stabilized microtubules (magenta) following incubation with 1 nM GFP-SEPT9_i1 (green) for wild type, and S82A/S85A, and S82E/S85E mutants. Scale bars, 10 μm. **(B)** Quantification of mean GFP fluorescence intensity per microtubule length following incubation with 1 nM GFP-SEPT9_i1 (*n* = 30; WT, S82A/S85A, S82E/S85E). Pairwise comparisons were statistically analyzed using an unpaired Welch’s t-test. *, p < 0.05; ****, p < 0.0001 **(C)** TIRF microscopy images of taxol-stabilized microtubules (magenta) following incubation with 1 nM of the indicated msGFP-tagged wild-type SEPT2/6/7/9_i1, SEPT2/6/7/9_i1-S82A/S82A, and msGFP-SEPT2/6/7/9_i1-S82E/S85E (green). Scale bars, 10 μm. **(D)** Quantification of mean GFP fluorescence intensity per microtubule length following incubation with 1 nM wild-type GFP-SEPT9_i1 (*n* = 30), msGFP-SEPT2/6/7/9_i1-S82A/S82A (*n* = 30), and msGFP-SEPT2/6/7/9_i1-S82E/S85E (*n* = 30). Pairwise comparisons were statistically analyzed using a Mann-Whitney U-test. ****, p < 0.0001 All plots show mean ± SEM.

### S82/S85 phosphomimetic mutants enhance actin binding of SEPT9_i1 and SEPT2/6/7/9_i1 complexes in vitro

To test whether phosphomutations at S82 and S85 SEPT9 affect actin filament binding directly, we generated phalloidin-stabilized actin filaments from rabbit skeletal muscle G-actin and immobilized them in glass chambers for TIRF microscopy-based binding assays. Wild type GFP-SEPT9_i1 decorated actin filaments at concentrations of approximately 100 nM (Figure 5A). Strikingly, and in contrast to the drastic reduction in microtubule-binding, the phosphomimetic S82E/S85E mutant increased actin-bound GFP-SEPT9_i1 by more than two-fold at 100 and 200 nM (Figure 1A-B and S6A-B). The phosphonull mutations S82A/S85A mutant, however, had no significant effect on actin decoration at either concentration (Figure 1A-B and S6A-B).

**Figure 5.**
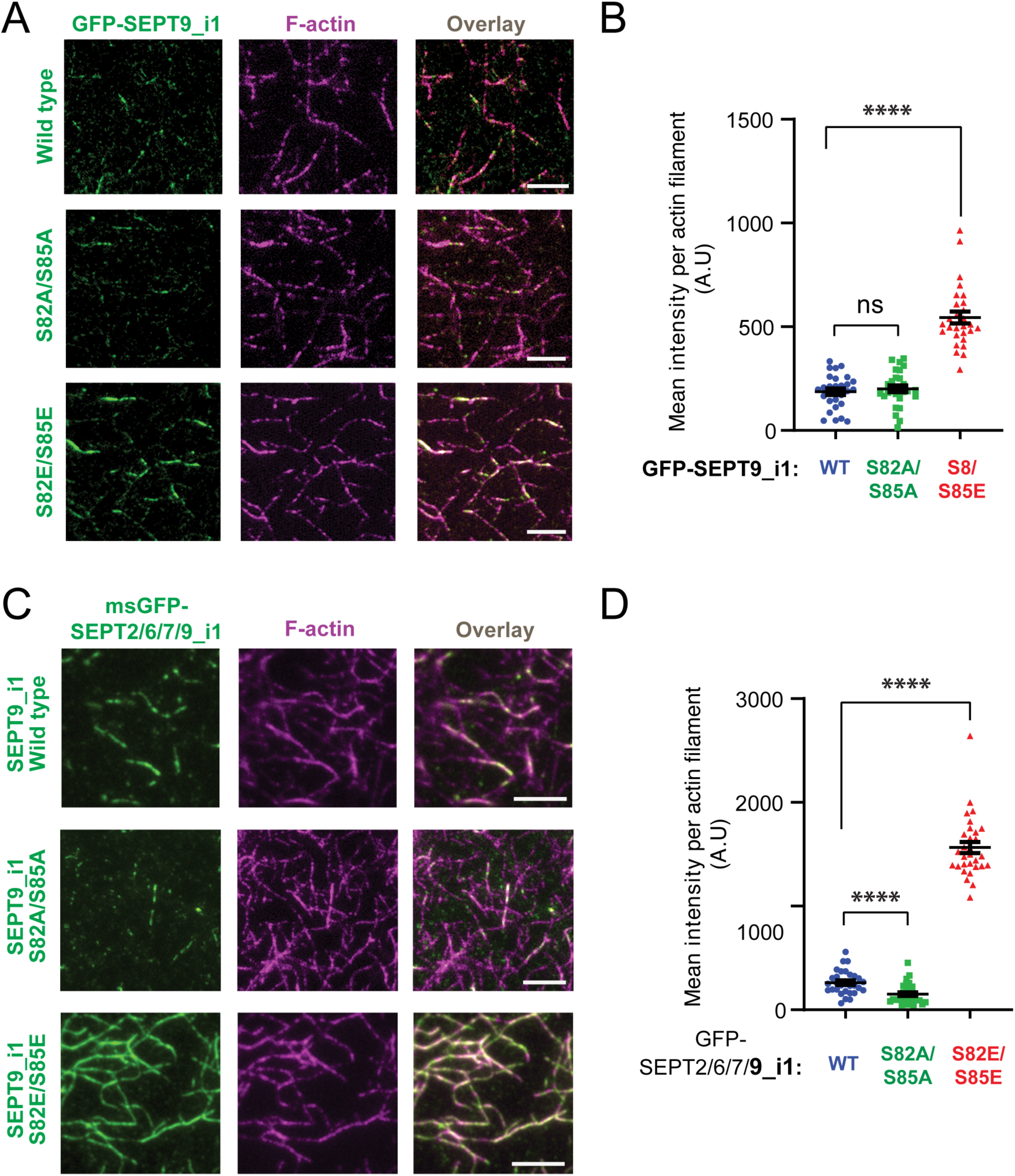
Phosphomimetic mutations at S82/S85 of SEPT9_i1 enhance actin filament binding in vitro. **(A)** Total internal reflection fluorescence (TIRF) microscopy images of phalloidin-stabilized actin filaments (magenta) following incubation with 100 nM GFP-SEPT9_i1 (green) for wild type, and the S82A/S85A and S82E/S85E mutants. Scale bars, 5 μm. **(B)** Quantification of mean GFP fluorescence intensity per actin length following incubation with 100 nM GFP-SEPT9_i1 (*n* = 28; WT, S82A/S85A, S82E/S85E). Pairwise comparisons between WT and S82A/S85A was statistically analyzed with a Welch’s t-test, and between WT and S82E/S85E with a Mann Whitney U-test. ns, not significant; ****, p < 0.0001 **(C)** TIRF microscopy images of phalloidin-stabilized actin filaments (magenta) following incubation with 1 μM of the indicated msGFP-tagged wild-type SEPT2/6/7/9_i1, SEPT2/6/7/9_i1-S82A/S82A, and msGFP-SEPT2/6/7/9_i1-S82E/S85E (green). Scale bars, 5 μm. **(D)** Quantification of mean GFP fluorescence intensity per actin length following incubation with 1 μM wild-type GFP-SEPT9_i1 (*n* = 30), msGFP-SEPT2/6/7/9_i1-S82A/S82A (*n* = 30), and msGFP-SEPT2/6/7/9_i1-S82E/S85E (*n* = 30). Pairwise comparisons were statistically analyzed using a Mann-Whitney U-test. ****, p < 0.0001 All plots show mean ± SEM.

To corroborate these findings with an independent approach, we assessed actin filament binding by incubating GFP-SEPT9_i1 in solution with phalloidin-stabilized actin filaments, which were pre-polymerized from human platelet G-actin, and imaging the resulting septin-decorated actin by TIRF microscopy. In this assay, and as previously reported ^20,74^, SEPT9_i1 bundles actin filaments. We, therefore, imaged the septin-decorated actin bundles after flowing into TIRF microscopy chambers. Consistent with enhanced single actin filament decoration, S82E/S85E increased both the number and length of actin bundles (Figure S6C-E), with the increase in bundle length suggesting a longitudinal staggering mechanism of filament crosslinking. In contrast, S82A/S85A decreased bundle formation but without affecting bundle length or single filament decoration (Figure S6C-E), suggesting that the phosphonull mutation impacts the efficiency of bundling rather than the underlying mechanism.

As SEPT9 isoforms are thought to drive the interaction of heteromeric septin complexes with actin filaments, we asked whether the phosphomutations affect actin-binding in the context of the hetero-octameric complex. The S82E/S85E mutant had an even more pronounced effect on GFP-SEPT2/6/7/9_i1 than GFP-SEPT9_i1 alone, increasing actin-bound complex by nearly seven-fold (Figure 5C-D). Conversely, S82A/S85A markedly reduced actin binding of the hetero-octamer (Figure 5C-D) – an effect that parallels the reduced bunding efficiency seen with the GFP-SEPT9_i1 mutant (Figure S6D), but contrasts with its lack of effect on single filament decoration (Figure 5B). This suggests that the impact of S82A/S85A is context dependent.

Within the hetero-octamer, where SEPT9 is the sole actin-binding subunit present once every four subunits, even modest reduction in individual binding affinity might be amplified, or the mutation may allosterically alter the actin-binding interface of the complex. Regardless of the context-dependent effects of the phosphonull mutant, the data collectively demonstrate that the phosphomimetic S82E/S85E strongly enhances actin binding – an effect strikingly opposite to its inhibitory effect on microtubule binding. These findings support a model in which GSK3-dependent phosphorylation of SEPT9_i1 at S82/S85 acts as a molecular switch, shifting the distribution of SEPT9-containing septin complexes between microtubules and actin filaments.

### Dephosphorylation of SEPT9_i1 at S82/S85 by GSK3β inactivation promotes microtubule association and the establishment of neuronal polarity

To examine the physiological significance of GSK3-mediated phosphorylation of SEPT9_i1 at S82/S85, we turned to neuronal morphogenesis, where GSK3 signaling plays a well-established role in controlling neuronal polarity ^59^. The establishment of polarity requires the asymmetric outgrowth of a single neurite that becomes the axon — a process driven by convergent signaling pathways that inactivate GSK3, thereby stabilizing microtubules through reduced phosphorylation of microtubule-associated proteins (MAPs) ^56,59^. Because SEPT9_i1 association with microtubules promotes asymmetric neurite growth^19^, we investigated whether GSK3 phosphorylation regulates SEPT9_i1 partitioning between the actin and microtubule networks in polarizing neurons, and whether this regulation contributes to the establishment of neurite asymmetry.

Using embryonic rat primary hippocampal neurons — an established *in vitro* model of neuronal polarization ^81^ — we characterized the steady-state localization of endogenous SEPT9 relative to actin filaments and microtubules. SEPT9 colocalized preferentially with microtubules, with the highest colocalization observed along the neurite shaft (Figure 6A-B). Pearson’s coefficients for colocalization with actin filaments of the axon shaft and growth cone were markedly lower (Figure 6C), confirming an overall preference for the microtubule network. Pharmacological inhibition of GSK3 with CHIR99021 further enhanced SEPT9 colocalization with microtubules in both the neurite shaft and the central domain of the growth cone (Figure 6A-B). Despite the low baseline colocalization with actin, CHIR99021 treatment also resulted in a statistically significant decrease in SEPT9 association with the growth cone actin network (Figure 6C), consistent with our observations in U2OS cells (Figure 1E).

**Figure 6.**
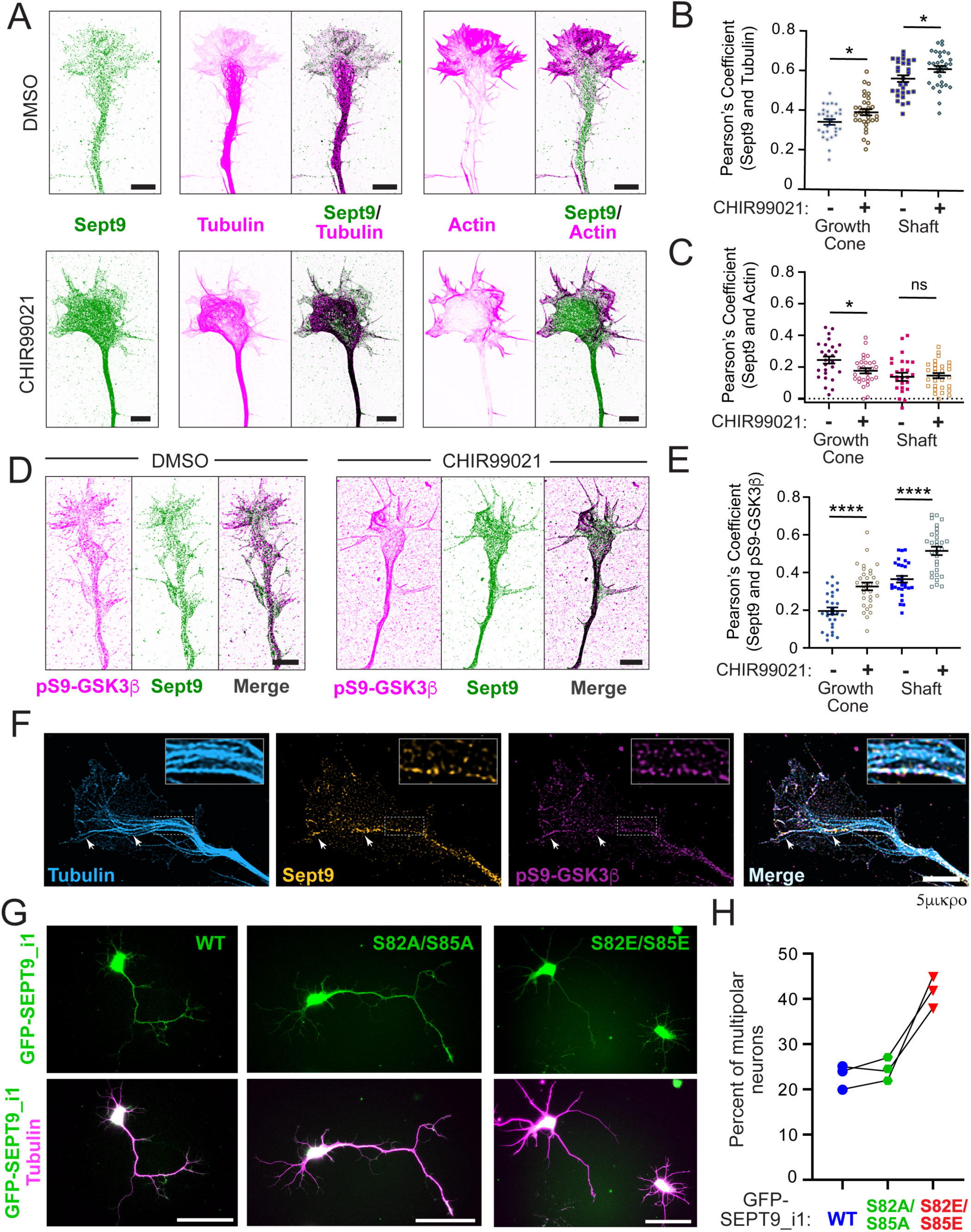
SEPT9 association with neuronal microtubules promotes asymmetric neurite growth in a GSK3 inactivation-dependent manner. **(A)** Super-resolution SoRa spinning-disk confocal microscopy images of growth cones in primary rat embryonic (E18) hippocampal neurons at day in vitro 2 (DIV2), following 1 h treatment with CHIR99021 (3 μM) or DMSO (control). Neurons were stained for endogenous Sept9 (inverted forest green), acetylated tubulin (inverted magenta) and filamentous actin (phalloidin; inverted magenta). Overlap between forest green and magenta colors appears black. Scale bars, 5 μm. **(B)** Pearson’s correlation coefficient (mean ± SEM) between Sept9 and acetylated tubulin fluorescent channels measured in growth cones and neurite shafts of the longest neurites (*n* = 29-32 neurons per condition) following 1 h treatment with CHIR99021 (3 μM) or DMSO. Pairwise comparisons were performed using an unpaired Welch’s t-test. *, p < 0.05 **(C)** Pearson’s correlation coefficient (mean ± SEM) between the Sept9 and phalloidin fluorescent channels in growth cones and neurite shafts of the longest neurites (*n* = 27 neurons per condition) following 1 h treatment with CHIR99021 (3 μM) or DMSO. Pairwise comparisons were performed using an unpaired Welch’s t-test. *, p < 0.05; ns, not significant **(D-E)** Super-resolution SoRa spinning-disk confocal microscopy images (D) of neurite growth cones in primary rat embryonic (E18) hippocampal neurons at DIV2, following 1 h treatment with CHIR99021 (3 μM) or DMSO. Neurons were co-stained for endogenous Sept9 and inactive GSK3β using a phospho-specific antibody against pSer9-GSK3β. Scale bars, 5 μm. Quantification **(E)** shows the Pearson’s correlation coefficients (mean ± SEM) between Sept9 and pSer9-GSK3β fluorescent channels in growth cones and shafts of neurites (*n* = 28-30 neurons per condition) for the indicated treatments. Pairwise comparisons were performed using an unpaired Welch’s t-test. ****, p < 0.0001 **(F)** Super-resolution SoRa spinning-disk confocal microscopy image of the growth cone of the longest neurite in a primary rat hippocampal neuron (DIV2), co-stained for endogenous acetylated tubulin (cyan), Sept9 (orange) and inactive pS9-GSK3β (magenta). Arrows indicate colocalization of Sept9 and pS9-GSK3β on distinct microtubule segments and tracks of the growth cone. Insets show the at higher magnification the region outlined by a dashed rectangle. Scale bars, 5 μm. **(G-H)** Wide field fluorescence microscopy images (G) of primary rat hippocampal neurons at DIV3, stained for α-tubulin (magenta), following co-transfection with plasmid expressing SEPT9 shRNA and shRNA-resistant GFP-SEPT9_i1 constructs (wild-type, S82A/S85A, or S82E/S85E). Scale bars, 50 μm. Quantification (H) shows the percentage of multipolar neurons – defined as neurons lacking single dominant asymmetric neurite – after 3 days of expressing wild type GFP-SEPT9_i1 (*n* = 33, 24, 35), GFP-SEPT9_i1-S82A/S85A (*n* = 52, 25, 32) and GFP-SEPT9_i1-S82E/S85E (*n* = 36, 21, 33). Results from three independent experiments are shown.

We next reasoned that if SEPT9 microtubule association is promoted by GSK3 inactivation, SEPT9 might be spatially enriched at microtubule segments decorated with inactive GSK3. To test this, we immunolabeled neurons with a phospho-specific antibody against pSer9-GSK3β — a well-characterized inhibitory phosphorylation mark ^62^ — and antibodies against SEPT9 and tubulin, and imaged cells by super-resolution SoRa spinning disk confocal microscopy. We analyzed SEPT9 and pSer9-GSK3β colocalization in mock and CHIR99021 treated neurons (Figure 6D). In mock-treated neurons, SEPT9 colocalization with pSer9-GSK3β was the highest in microtubule-rich neurite shafts, and inhibition of GSK3β activity by CHIR99021 increased SEPT9 colocalization with pSer9-GSK3β in both neurite shafts and growth cones (Figure 6D-E). These data suggested that local GSK3β inactivation promotes SEPT9 association with neuronal microtubules, raising the possibility that SEPT9 is enriched on microtubule segments and/or microtubules subsets with inactive GSK3β. Indeed, super-resolution SoRa imaging showed that SEPT9 is co-enriched with pSer9-GSK3β domains on distinct microtubule segments and microtubules, which extended from the neurite shaft into both the central and peripheral domains of the growth cone (Figure 6F).

To directly test whether GSK3β-mediated phosphorylation of SEPT9_i1 regulates neuronal polarization, we expressed the wild-type, phosphonull (S82A/S85A) and phosphomimetic (S82E/S85E) mutants of GFP-SEPT9_i1 while depleting endogenous SEPT9 in rat hippocampal neurons at DIV0, and assessed neurite asymmetry at DIV3. Neurons expressing wild-type or phosphonull GFP-SEPT9_i1 developed robust neurite asymmetry at comparable rates, with approximately 80% exhibiting a single dominant neurite at least twice the length of the next shortest — consistent with a presumptive axon — alongside a characteristic pyramidal morphology with shorter neurites emanating from the opposite side of the cell body (Figure 6G). In contrast, expression of the phosphomimetic S82E/S85E mutant, which has reduced microtubule association, impaired neurite asymmetry, resulting in an approximately two-fold increase in multipolar neurons — neurons that possess multiple neurites of similar length and lack a dominant outgrowth (Figure 6G-H). Collectively, these results demonstrate that SEPT9_i1 promotes neuronal polarization in its unphosphorylated form, and that GSK3β inactivation — by relieving phosphorylation at S82/S85 — enables SEPT9_i1-dependent microtubule stabilization that drives the asymmetric neurite outgrowth required for the establishment of neuronal polarity.

## Discussion

Septins comprise a distinct network of cytoskeletal filaments that interact with actin filamants and microtubules, and spatially regulate cellular processes driven by or dependent on these cytoskeletal networks ^2^. Intriguingly, septins are less abundant than actin filaments and microtubules in both absolute protein concentration and polymer mass, raising a fundamental question: how do septin complexes sort between actin and microtubule subnetworks, and what mechanisms regulate these interactions? This knowledge gap has been further compounded by a poor understanding of the biochemical and structural basis of septin interactions with these cytoskeletal networks.

Here, we have identified a GSK3-dependent phosphorylation switch as a key modulator of septin portioning between microtubules and actin filaments, linking septin cytoskeletal functions to signaling pathways that govern cellular morphogenesis, metabolism and homeostasis. GSK3-mediated phosphorylation of SEPT9 at S82/S85 represents the first post-translational modification shown to bidirectionally gate septin-microtubule and septin-actin interactions — promoting the latter while inhibiting the former. We envisage that this mechanism operates in parallel with the Rho GTPase Cdc42 and its Cdc42EP effector proteins, which are required for binding to actin ^26,40^. Although phosphorylation of septins by various kinases has previously been reported to alter septin localization and protein interactions ^54,55^, this is the first phosphorylation event demonstrated to bidirectionally control septin interactions with both microtubules and actin.

The inhibitory effect of S82/S85 phosphorylation on microtubule binding is consistent with the paradigm of electrostatic regulation of MAP-microtubule interactions, which initially emerged in the context of tau hyperphosphorylation in neurodegeneration ^52,82,83^. In this paradigm, phosphorylation of microtubule-binding domains — typically within repeat motifs and adjacent proline-rich regions — introduces negative charge that clashes with the acidic C-terminal tails of tubulin, progressively reducing binding affinity ^52,84,85^. S82 and S85 reside within a ∼60 amino acid basic domain that is itself sufficient to directly bind and bundle microtubules *in vitro*, flanked by two clusters of K/R-X-X-D/E motifs ^19^. The basic residues within these motifs mediate heterophilic electrostatic interactions with the acidic C-terminal tails of tubulin, as well as homophilic interactions with the D/E residues of neighboring motifs that crosslink septin-bound microtubules ^19^. Phosphorylation at S82/S85 is therefore predicted to disrupt both classes of electrostatic interaction, simultaneously impairing microtubule binding and bundling.

Whether the first ∼25 amino acids of the SEPT9 N-terminus — which harbor MAP-like microtubule-binding motifs ^22^ — are independently regulated by phosphorylation at their four serine and one threonine residues remains to be determined. These residues are spatially distal to S82/S85, but if the intrinsically disordered N-terminal domain adopts a more compact conformation upon microtubule association, S82/S85 phosphorylation could influence this region. The basic domain harboring S82/S85 is not itself enriched in prolines, which are more abundant in the neighboring acidic domain. Given that proline-rich sequences can adopt polyproline II (PPII) helix conformations ^86^ — extended, three-residue-per-turn helices that periodically expose basic residues for interaction with the acidic C-terminal tails of tubulin — S82/S85 phosphorylation may additionally interfere with PPII-mediated tubulin contacts, either electrostatically or allosterically. Whether PPII helices form in the N-terminus of SEPT9 remains uncertain, however, as this region is predicted to be intrinsically disordered, and structural data for this domain in the context of microtubule binding are currently lacking.

In striking contrast to its effect on microtubule binding, S82/S85 phosphorylation enhances binding to actin filaments. This phosphorylation-dependent gain of actin affinity parallels observations in the ERM family (ezrin, radixin, moesin), where phosphorylation promotes actin binding ^87,88^, and in cortactin, where Src- and ERK-mediated phosphorylation increases actin nucleation activity ^89,90^. However, the underlying mechanism in SEPT9 is likely distinct. ERM proteins are activated indirectly — phosphorylation relieves intramolecular autoinhibition rather than directly remodeling the actin-binding interface ^91,92^. For SEPT9, prior work has identified three alternate docking modes by which full-length SEPT9 and its basic domain associate with actin filaments, each engaging distinct filament surfaces; only one partially overlaps with the ERM actin-binding surface ^20^. How phosphorylation at S82/S85 mechanistically may favor one or more of these binding modes is unclear.

The regulatory complexity of the SEPT9 N-terminal domain extends beyond S82/S85. Phosphorylation of S30 — a residue conserved across SEPT9 isoforms 1, 2, and 3 — has been indirectly implicated in actin binding in the context of enteropathogenic *E. coli* (EPEC) infection, where the bacterial type III secretion system induces S30 phosphorylation to promote SEPT9 assembly with actin at sites of bacterial attachment ^93^. Together with our findings, the S30 data raise the possibility that the SEPT9 N-terminal domain harbors multiple phosphorylation-sensitive nodes that independently or cooperatively tune actin binding in response to distinct upstream signals. CDK1-mediated phosphorylation of T24 in SEPT9 isoform 3 — a threonine conserved across the three longest SEPT9 isoforms — regulates SEPT9 localization at the intracellular bridge during the abscission phase of cytokinesis ^94^, though whether this phosphorylation event impacts actin or microtubule binding is unknown. Similarly, S93F in SEPT9 isoform 3 – also conserved across the longest SEPT9 isoforms - is clinically linked to the pathogenesis of the HNA neuropathy ^95,96^, but it is unknown if it impacts microtubule and actin binding. Beyond phosphorylation, the N-terminal domain of SEPT9 has been identified as a site of ubiquitination and sumoylation ^97,98^, which points to a level of regulatory complexity that reflects the unique structural and functional versatility of the SEPT9 N-terminal extension, which may integrate multiple post-translational signals to tune septin interactions with the cytoskeleton.

In most tissues, GSK3 is constitutively active and its inhibition through phosphorylation by upstream signaling kinases – e.g., Akt and the insulin signaling pathway – produces downstream effects by remodeling key molecular interactions ^58,59,62^. In the case of SEPT9, constitutive GSK3 activity would maintain S82/S85 phosphorylation at steady state, biasing septin distribution toward the actin cytoskeleton. This is consistent with the well-established predominance of septin association with actin stress fibers and cortical actin as the default localization pattern in most mammalian cell types ^2,5^. Notably, however, the priming serine at the +4 position of the consensus GSK3β site is absent in some mammals, suggesting that S85 might not be constitutively phosphorylated in all species. In such organisms, the longest SEPT9 isoforms might more extensively associated with microtubules under basal conditions. This species-level variation raises the broader possibility that the relative weighting of septin distribution between actin and microtubules is not fixed but has been tuned evolutionarily through changes in the priming phosphorylation context.

The transition from actin- to microtubule-associated SEPT9 would require not only GSK3 inhibition but also active dephosphorylation of pS85 by as yet unidentified phosphatases. This is likely to be a kinetically constrained process. Because S85 is a proline-directed phosphosite, the flanking proline is expected to sterically impede phosphatase access, necessitating prior isomerization of the phospho-Ser/Pro peptide bond by the prolyl isomerase Pin1 before dephosphorylation can occur — a regulatory mechanism previously proposed for the CDK1-dependent T24 phosphorylation site of SEPT9 isoform 3 ^63,64,94^. This multi-step dephosphorylation requirement would impose a significant kinetic barrier on the actin-to-microtubule transition, effectively functioning as a molecular delay that prevents spurious or transient fluctuations in GSK3 activity from immediately redistributing the septin network. This may also explain why pharmacological GSK3 inhibition in this study did not lead to wholesale redistribution of SEPT9 from actin to microtubules. Taken together, these considerations suggest that septin association with microtubules is subject to a layered and temporally controlled regulatory mechanism — one that requires the coordinated action of GSK3 inhibition, Pin1-mediated prolyl isomerization, and phosphatase recognition — conferring robustness and directionality on what might otherwise be a reversible and noisy phosphorylation switch.

In conclusion, the identification of GSK3-dependent control of SEPT9 cytoskeletal partitioning provides a new conceptual framework for understanding how septin organization is coupled to upstream signaling in both physiological and disease contexts. In neuronal morphogenesis, our findings support a model in which GSK3β inactivation promotes SEPT9 association with microtubules, which in turn drives the asymmetric neurite outgrowth that underlies axon specification. Beyond neuronal polarity, GSK3 inhibition by insulin signaling may regulate SEPT9-microtubule association in the context of glucose transporter vesicle trafficking and lipid droplet biogenesis — processes in which both septins and microtubules are implicated ^28,99^. In cancer, where GSK3β activity can exert both tumor-suppressive and oncogenic functions in a context-dependent manner ^100,101^, alterations in the balance of SEPT9 localization between actin and microtubules may contribute to or reinforce mechanisms of tumor suppression or progression. Future work aimed at dissecting the functional and mechanistic intersections between SEPT9, GSK3, and the signaling networks that regulate them will be essential for understanding the pathophysiology of diseases in which both proteins are dysregulated, and for evaluating their potential as therapeutic targets.

### Limitations of the Study

Pharmacological inhibition of GSK3 activity impacted the localization of endogenous SEPT9 in multiple cell types, but this effect was variable and relied on the use of the CHIR99021 compound. Future work will investigate whether GSK3 targets additional residues and sites in the N-terminal microtubule- and actin-binding domains of SEPT9, and the role of other kinases in priming the GSK3 phosphorylation events as well as the role of proline isomerases such as Pin1 in further regulating the phosphorylation and dephosphorylation cycle of S85. Future work will also explore the structural basis of the effects of SEPT9_i1 phosphorylation on microtubule and actin binding, and how phoshosphorylation may impact septin-mediated microtubule-actin crosstalk.

## RESOURCE AVAILABILITY

### Lead Contact

Requests for further information and resources should be directed and will be fulfillted by the lead cotact, Elias T. Spiliotis

### Materials Availability

All materials used in this study will be made available upon request to the lead contact and after the completion of material transfer agreements, if necessary.

## ACKNOWLEDGMENTS

This work was supported with a National Institute of General Medical Sciences (NIGMS) grant R35GM136337 from the National Institutes of Health (NIH) to E.T.S.

## AUTHOR CONTRIBUTIONS

Conceptualization, M.N.A.A., A.W.S., P.B., I.K., and E.T.S.; investigation, M.N.A.A., T.C.H., A.W.S., S.R., F. M., S.B., P.B., I.K., and E.T.S.,; methodology, M.N.A.A., T.C.H., A.W.S., S.R., F. M., S.B., P.B., I.K., and E.T.S.; supervision, E.T.S.; writing – original draft, M.N.A.A., J.C.H., A.W.S., S.R., F.M., and E.T.S.; writing – review & editing, M.N.A.A. and E.T.S.

## STAR METHODS

### EXPERIMENTAL MODEL AND STUDY PARTICIPANT DETAILS

#### Cell lines and cell culture

U2OS cells (ATCC HTB-96) were maintained in DME High Glucose medium (Sigma-Aldrich-07777). MDCK cells (RRID:CVCL_0424) cells were maintained in low glucose DME (Cat# D5523; Sigma Aldrich) with 1 g/L NaHCO3 (Cat# S5761; Sigma-Aldrich). All media were supplemented with 10% fetal bovine serum (Atlas Biologicals FR-0500-A) and 1% penicillin (P3032; Sigma-Aldrich), streptomycin (Cat# 11860-038; Gibco), and kanamycin (Cat# K4000; Sigma-Aldrich) at 37°C in a 5% CO2 incubator. Cells were maintained in culture at 37 °C in a humidified incubator with 5% CO₂. Cultures were maintained at 60%-70% confluence and sub-cultured at a 1:5–1:10 ratio every 4 days using 0.05% trypsin-EDTA (Gibco, 25300-054). Prior to experiments, cells were seeded on glass coverslips of #1.5 thickness (Cat# 48366227; VWR), which were coated with type I collagen (PureCol type I bovine collagen; Advanced Biomatrix, 5005). Glass coverslips were coated with collagen at a concentration of 40 µg/mL for 2 minutes at room temperature, and subsequently were air-dried for 30 minutes and exposed to UV light for 30 minutes in a biosafety cabinet to ensure sterilization. For experiments performed on glass coverslips, cells were detached using 0.05% trypsin-EDTA for 5 minutes at 37 °C, neutralized with complete DMEM (DMEM High Glucose medium, 10% fetal bovine serum and 1% PSK), and collected by centrifugation at 1,000 rpm for 3 minutes. The cell pellet was resuspended in fresh complete DMEM, and viable cells were counted using an automated cell counter Countess 3 (Thermo Fisher Scientific) with 0.4% (w/v) trypan blue exclusion (Corning, 25-900-Cl). Cells were seeded at a density of 2 × 10⁵ cells per coverslip and allowed to adhere and recover for 12–18 hours before transfection.

Primary neurons were dissociated from freshly dissected embryonic (E18) rat hippocampi (Transnetyx Tissue). Hippocampi were incubated for 10 minutes at 34-36 °C in Hibernate E medium without calcium chloride (Transnetyx Tissue) supplemented with 2 mg/mL papain, and triturated 10 times or until homogeneous using a fire-polished long-stem Pasteur glass pipette (Corning). Cells were pelleted, counted, and plated at a density of 150,000 cells per 22 mm glass coverslip (No. 1.5; VWR). Glass coverslips were coated with 0.1 mg/mL poly-D-lysine (Sigma-Aldrich P0899) overnight at 37°C followed by 1.25 µg/mL laminin (R&D Systems #3400-010-02) for 1 hour at 37°C.

In experiments that involved GSK3 inhibition, U2OS and MDCK cells were seeded at a density of 2.5 × 10⁵ per 35-mm dish and incubated with CHIR-99021 (Cell Signaling # 54290; 10 μM) or equal volume vehicle control (DMSO) in regular DME media for 2-12 hour at 37°C. Primary rat hippocampal neurons were treated with the CHIR-99021 (3 µM) or vehicle (DMSO) control for 1 hour at 37°C.

In experiments of microtubule depolymerization and regrowth, MDCK cells were seeded at a density of 2 × 10⁵ cells per 35-mm dish with 22×22 mm collagen-coated glass coverslips. Forty-eight hours after transfection, microtubules were depolymerized by treating cells with 1.6 µM nocodazole (Sigma-Aldrich # M1404) for 2 h at 37°C, and were allowed to regrow by rinsing once with and then incubating for 40 seconds in fresh, pre-warmed nocodazole-free culture medium to allow microtubule regrowth prior to fixation.

### METHOD DETAILS

#### Mutagenenesis

Site-directed mutagenesis was performed to generate phosphonull (S82A/S85A) and phosphomimetic (S82E/S85E) His-GFP-SEPT9_i1 variants using Gibson assembly. Mutations were introduced into the pET-28a(+) plasmid expressing His-GFP-SEPT9_i1 (GenBank accession number NP_001106963; isoform a), which has been described previously ^80^ using the following forward and reverse primers: 5′-GCCCGCCATGTGGACTCCCTATCACAACGCGCGCCCAAGGCGTCCCTG-3′ (S85A forward), 5’-CAGGGACGCCTTGGGCGCGCGTTGTGATAGGGAGTCCACATGGCGGGC-3’ (S85A reverse), 5′-GCCCGCCATGTGGACTCCCTATCACAACGCGAGCCCAAGGCGTCCCTG-3′ (S85E forward), 5’-CAGGGACGCCTT GGGCTCGCGTTGTGATAGGGAGTCCACATGGCGGGC-3’ (S85E reverse), 5′-GCCCGCCAT GTGGACTCCCTAGCGCAACGCGCGCCCAAGGCGTCCCTG-3′ (S82A/S85A forward), 5’-CAGGGACGCCTTGGGCGCGCGTTGCGCTAGGGAGTCCACATGGCGGGC-3’ (S82A/S85A reverse), 5′-GCCCGCCATGTGGACTCCCTAGAGCAACGCGAGCCCAAGGCGTCCCTG-3′ (S 82E/S85E forward), and 3’-CAGGGACGCCTTGGGCTCGCGTTGCTCTAGGGAGTCCACATGGCGGGC-5’ (S82E/S85E reverse). The same primers were used to mutate an shRNA-resistant GFP-tagged version of the human SEPT9 isoform 1 in a pSUPER plasmid that co-expressed the shRNA targeting sequence 5’-GCACGATATTGAGGAGAAA-3’ (1038) to deplete endogenous SEPT9, which was previously constructed and used ^68,102^. All constructs were verified by DNA sequencing prior to use.

For the recombinant SEPT9_i1 expressed in bacteria, site-directed mutagenesis was performed to generate phosphonull (S82A/S85A) and phosphomimetic (S82E/S85E) His-GFP-SEPT9_i1 variants via Gibson assembly using the primers above and the plasmid His-GFP-SEPT9_i1 (GenBank accession number NP_001106963; isoform a), which was created by PCR amplifying GFP-SEPT9 and inserting it into the pET-28a(+) vector, and has been described previously^80^.

For the SEPT2/6/7/9_i1 complex, we used the plasmid pnCS_SEPT7_SEPT9_i1-TEV-Strep (Addgene plasmid 174500; http://n2t.net/addgene:174500; RRID:Addgene_174500), and the plasmid pnEA-vH_His-TEV-SEPT2-msfGFP_SEPT6 (Addgene plasmid # 174498; http://n2t.net/addgene:174498; RRID:Addgene_174498), both of which were gifts from Dr. Manos Mavrakis ^78^. The Phospho-mimetic and phospho-null mutant of Septin9 in the Septin2/6/7/9_i1 octamer was generated using the Q5 Site directed mutagenesis (SDM) kit (E0554S; New England Biolabs). Site directed mutagenesis was performed to introduce a substitution mutation into the SEPT9_i1 gene in pnCS_SEPT7_SEPT9_i1-TEV-Strep (Addgene plasmid 174500; http://n2t.net/addgene:174500; RRID:Addgene_174500) to replace the S82 and S85 with A (alanine) and E (glutamate) for phospho-mimetic and phosphor-null mutants respectively. The following mutagenic primers were used 5’-cgcgcgCCCAAGGCGTCCCTG-3’ (S82A/85A forward), 5’-ttgcgcTAGGGAGTCCACATGGC-3’ (S82A/85A reverse), 5’-cgcgagCCCAAGGCGTCCCTG-3’ (S82E/S85E forward), 5’-ttgctcTAGGGAGTCCACATGGC-3’ (S82E/S85E reverse), which designed using the Online NEB base changer tool. All mutations were confirmed by Sanger sequencing by targeting the T7 promoter region.

#### Transfections

U2OS and MDCK cells were transfected transiently with plasmids using Lipofectamine 2000 (Invitrogen, 11668019) according to the manufacturer’s instructions with minor optimization. Transfections were carried out in culture dishes using Opti-MEM™ I Reduced Serum Medium (Gibco, Cat. #31985070). Plasmid DNA–lipid complexes were freshly prepared immediately prior to transfection. Unless otherwise specified, 500 ng of plasmid DNA for MDCK cells or 1 µg of plasmid DNA for U2OS cells were diluted in 100 µL of Opti-MEM™ I. In a separate tube, 4 µL of Lipofectamine 2000 was diluted in 100 µL of Opti-MEM™ I. Both solutions were incubated at room temperature for 5 minutes before being combined. The mixtures were gently mixed and allowed to incubate for an additional 20 minutes at room temperature to allow formation of DNA–lipid complexes. Prior to transfection, the existing culture medium was replaced with fresh DMEM Glucose medium lacking antibiotics. The prepared DNA–Lipofectamine 2000 complexes were then added dropwise to the cells, and the dishes were gently swirled to ensure even distribution. Cells were maintained at 37 °C in a humidified incubator with 5% CO₂ for 6 hours. Following this incubation period, the transfection medium was replaced with complete DMEM with antibiotics, and the cells were allowed to continue expressing the constructs for a total of 48 hours in MDCK cells or 24 hours in U2OS cells. Primary neurons were electroporated immediately prior to plating using Nucleofector II (Amaxa) with Ingenio Electroporation Kits and Solution (Mirus Bio). Cells were maintained in Neurobasal Medium with 1X B27 and 1X GlutaMAX supplements (ThermoFisher Scientific) and incubated at 37°C with 5% CO2.

#### Immunofluorescence

U2OS and MDCK were fixed with 4% paraformaldehyde (Electron Microscopy Sciences, Cat. #15710) prepared in PHEM (60 mM PIPES, 25 mM HEPES, 10 mM EGTA, and 2 mM MgCl₂ pH 6.9) for 20 min at room. In Figures 1F, 3A, S2A and S3, cells were pre-extracted for 10 s using PHEM buffer containing 0.5% Triton X-100 and 4% paraformaldehyde (PFA) to remove soluble tubulin and cytosolic proteins while preserving polymerized microtubules prior to fixation and staining. To neutralize residual aldehyde groups generated during fixation, the cells were treated with 0.4% ammonium chloride (Sigma, Cat. #A9434) for 5 minutes at room temperature. Cells were then permeabilized using 0.2% Triton X-100 (Sigma, Cat. #X100) in PBS for 20 minutes at room temperature. Following permeabilization, nonspecific binding sites were blocked with a blocking solution consisting of 2% bovine serum albumin (BSA; Sigma, Cat. #A9647) and 0.1–0.2% Triton X-100 in PBS for 30 minutes. The samples were subsequently rinsed twice with PBS prior to antibody staining. Primary antibodies diluted in blocking buffer were applied to the coverslips and incubated overnight at 4 °C. After primary antibody incubation, the coverslips were washed with PBS and incubated with the appropriate fluorophore-conjugated secondary antibodies for 1 hour at room temperature in the dark. After completion of the immunostaining procedure, the coverslips were washed with PBS and mounted onto frosted microscope slides (Globe Scientific, Cat. #1304W) using FluorSave™ mounting reagent (Millipore, Cat. #345789). Prepared slides were stored at 4 °C and protected from light until imaging.

Rat hippocampal neurons are pre-extracted to preserve the cytoskeleton using the following pe-extraction buffer for 20 seconds: 80 mM K-PIPES pH 6.9, 1mM EGTA, 150 mM NaCl, 7 mM MgCl2, 5 mM D-Glucose, 0.125% glutaraldehyde, 0.1% Triton X-100. Upon the pre-extraction, rat hippocampal neurons were fixed for 15 minutes in phosphate buffered saline (PBS) containing 4% paraformaldehyde (PFA; Electron Microscopy Sciences) and 4% sucrose. Cells were then simultaneously permeabilized and blocked with 5% bovine serum albumin (BSA), 0.3% saponin, and 0.2% sodium azide for 30 minutes. Primary antibodies were diluted in blocking solution and incubated at 4°C overnight or room temperature for 2 hours. Secondary antibodies and phalloidin were also diluted in blocking solution and incubated at room temperature for 2 hours. Coverslips were then mounted with FluorSave Reagent (MilliporeSigma).

#### Antibodies and Fluorescent Probes

The following primary antibodies were used for immunofluorescence microscopy and western blots: rabbit anti-Septin-9 (1:300, Proteintech, 10769), rabbit anti-Septin-9 (1:200, Milipore Sigma, HPA042564), anti-α-tubulin (1:300, Sigma-Aldrich, T6199), mouse anti-6XHIS (1:3000, BD Biosciences, 552565), rabbit anti-Septin-2 (1:1000, Kinoshita, 050830 aN5N(466) Rab 2g/L), ), rabbit anti-Septin-7 (1:400, IBL, 18991), mouse anti-α-tubulin (12G10, 1:1200; DSHB), rabbit Phospho-Serine/Threonine Polyclonal Antibody (1:3000, Bioss; bs-11994R-TR), mouse anti-phosphoserine/threonine monoclonal antibody (50-172-0851, 1:3000; ECM Biosciences), mouse Phospho-GSK3B (Ser9) monoclonal antibody (67558-1-Ig, 1:200; Proteintech), and mouse anti-acetylated tubulin antibody monoclonal (T7451, 1:1000; Sigma-Aldrich).

To minimize non-specific background staining in western blots with antibodies against phospho-serine/threonine, we precleared primary antibodies by adsorption with unphosphorylated recombinant SEPT9_i1 prior to use. We first coupled His-tagged GFP-SEPT9_i1 (3 µg) to Ni-NTA agarose resin (10–20 µL slurry; ∼5–10 µL settled resin; HisPur Ni-NTA Resin, Thermo Scientific # 88221) in binding buffer containing 50 mM Tris-HCl (pH 7.4–8.0), 300 mM NaCl, and 0.1% Triton X-100. Prior to binding, resin was equilibrated by washing 3 times with binding buffer following centrifugation (1,000×g, 1 min, 4°C). His–GFP-SEPT9_i1 was diluted in 200–300 µL binding buffer and incubated with the equilibrated resin for 2 hours at 4°C with gentle rotation. Following incubation, the resin was washed 3 times with binding buffer (1,000×g, 1 min, 4°C). The washed resin was then incubated with mouse phospho-threonine–proline monoclonal antibody (1:5,000; Cell Signaling Technology, #9391) in 300 µL binding buffer overnight at 4°C with rotation. After antibody incubation, samples were centrifuged (1,000xg, 1 min, 4°C), and the supernatant containing pre-cleared antibody was collected and for use in immunoblotting.

The following fluorescent secondary antibodies were used for immunodetection of primary antibodies in immunofluorescence experiments: Alexa Fluor 488 AffiniPure F(ab)₂ donkey anti-rabbit IgG (H+L) (1:200, Jackson ImmunoResearch, 712-545-153), AlexaFluor 488 AffiniPure F(ab)₂ donkey anti-mouse IgG (H+L) (1:200, Jackson ImmunoResearch, 715-546-150), Alexa Fluor 547 AffiniPure F(ab)₂ donkey anti-Rabbit IgG (H+L) (1:200, Jackson ImmunoResearch, 711-546-152), Alexa Fluor 488 anti-mouse IgM (1:1000, 406521, BioLegend). Actin was labeled with Phalloidin-iFluor 647 (Abcam, ab176759). The following secondary antibodies were used for detection of primary antibodies in western blots: IRDye 680LT goat anti-mouse IgM (μ chain specific; LICORbio 925-68080), IRDye 800 CW goat anti-mouse (LICORbio 926-32210), IRDye 800CW donkey anti-rabbit (LICORbio 926-32213), IRDye 680RD donkey anti-rabbit (LICORbio 926-68073) and IRDye 680 donkey anti-mouse (LICORbio 926-32222).

#### Protein expression and purification

Recombinant His-tagged GFP-SEPT9_i1, GFP-SEPT9_i1(S82A/S85A) and GFP-SEPT9_i1(S82E/S85E) were transformed into E.coli BL21 (DE3) (Invitrogen). Bacterial cultures were grown to OD600 of 0.5 and induced with 0.5 mM IPTG for 5 h at 22 °C or 0.2 mM IPTG for 16 h at 18 °C for fluorescence tagged SEPT9_i1. Cultures were centrifuged at 4,000 rpm for 20 min at 4 °C. Pellets were resuspended in lysis buffer containing 50 mM Tris pH 8.0, 150 mM NaCl, 10% glycerol, 1 mM PMSF, 1 mg/mL lysozyme, 10 mM imidazole and Bacterial Protease Arrest cocktail (G-Biosciences; 786–330) and lysed by sonication (8 rounds of pulsing for 30 sec followed by a 30 sec pause between each round). Cell lysates were centrifuged at 13,000 rpm for 30 min at 4 °C and passed through a 0.45 μm pore filter. Supernatants were loaded onto gravity flow columns with Ni-NTA agarose beads (745,400.25; Macherey-Nagel), which were equilibrated with 10 mL lysis buffer. Columns were washed with 30 mL washing buffer (50 mM Tris pH 8.0, 300 mM NaCl, 10% glycerol, 10 mM imidazole). Proteins were eluted in elution buffer containing 50 mM Tris pH 8.0, 150 mM NaCl, 10% glycerol and 250 mM imidazole, and dialyzed overnight in BRB80 (80 mM Pipes pH 6.9, 2 mM MgCl2, 1 mM EGTA).

Recombinant SEPT2/6/7/9 complexes (wild-type and mutants) were expressed in bacteria by co-transformaing with plasmids encoding SEPT2/6 and SEPT7/9 into E. coli BL21(DE3) strain, and growing LB plates with 100 μg/ml of Ampicillin and Spectinomycin and incubated at 37°C overnight. Single colony was picked and inoculated in LB containing 100 μg/mL of ampicillin and 50 μg/mL spectinomycin and incubated at 37°C until the culture reached an OD of 0.6-0.8. The cells were then induced with 0.5 mM IPTG for 16 h at 18°C. The induced cells were harvested by centrifugation at 4,000 RPM for 20 min at 4°C and resuspended in lysis buffer containing 100 mM Tris-HCl pH8, 150 mM NaCl, 1 mM EDTA and 1mM DTT, 1mM PMSF and 1 mg/ml DNase and lysed by sonication (10 Amplitude, 8 minutes, 30 seconds pulse on with 30 seconds interval). The lysed cells were then centrifuged at 20,000 RPM for 40 min and the supernatant was filtered through a 0.45 μm pore syringe filter. The purification was conducted on the AKTA pure FPLC system (Cytiva). The filtered lysate was applied on an equilibrated HisTrap HP 5 mL column (17524701; Cytiva) through a 50 ml Superloop(19785001; Cytiva). The HisTrap HP column was equilibrated using the binding buffer containing 100 mM Tris-HCl pH8, 150 mM NaCl, 1 mM EDTA and 1 mM DTT. After binding, the column was washed with wash buffer I (100 mM Tris-HCl pH 8, 150 mM NaCl, 1 mM EDTA,1 mM DTT and 15 mM Imidazole pH8) followed by a second wash with wash buffer II (100 mM Tris-HCl pH 8, 150 mM NaCl, 1 mM EDTA, 1 mM DTT and 40 mM Imidazole pH8). After the wash, the complex was eluted with an elution buffer (100mM Tris-HCl pH 8, 150 mM NaCl, 1 mM EDTA and 1 mM DTT and 250 mM Imidazole pH 8) at fixed fractions.

The eluted fractions were loaded onto a StrepTrap XT 1mL column (29401317; Cytiva) through a capillary loop. The StrepTrap XT column was equilibrated using the same binding buffer as with the HisTrap HP 5 mL. After sample application, the column was washed for five column volumes of binding buffer. The elution was conducted in two steps. The first elution step was set as two-step processes. In the first step, elution was performed with one column volume and subsequently paused for 30 min for equilibration with StrepTrap XT elution buffer (100 mM Tris-HCl pH 8, 150 mM NaCl, 1 mM EDTA, 1 mM DTT and 50 mM biotin). In the second step following the 30 min equilibration, elution proceeded with four column volumes of StrepTrap XT elution buffer. The fractions were collected based on the absorbance at 280 nm, and after pooling, they were further purified using a Superdex (SD) 200 Increase 10/300GL Gel filtration column (28990944; Cytiva) that was pre-equilibrated with buffer containing 50 mM Tris-HCl pH 8, 300 mM KCl, 5 mM MgCl2 and 3 mM DTT. Fractionation was performed with the same buffer. Fractions corresponding to the elution peak were pooled and snap-frozen for storage at -80°C after quantifying protein content and confirming protein identity by SDS-PAGE and western blotting.

#### In vitro GSK3β kinase assay

Recombinant GFP–SEPT9_i1 (wild-type, S82A/S85A, or S82E/S85E) (0.8 µg) was incubated with 1 µg recombinant human GSK3β (His-tagged; Sino Biological, 10044-H07B-50) in kinase reaction buffer containing Kinase Assay Buffer II (Sino Biological, K03-09-20), 1X PhosSTOP (Roche, 4906845001), 1 mM ATP, 1 mM PMSF, 1X bacterial protease inhibitor cocktail, and 0.25 mM DTT. Reactions were carried out at 30°C for 30 min or 4 h. Reactions were terminated by addition of SDS sample buffer (62.5 mM Tris-HCl (pH 6.8), 2% (w/v) SDS, 10% (v/v) glycerol, 0.01% (w/v) bromophenol blue, and 50 mM DTT) followed by denaturation at 95°C for 6 min. Protein samples were resolved on mPAGE 4–20% Bis-Tris precast gels (Millipore, MP42G15) at 200 V for ∼30 min. Proteins were transferred onto 0.45 µm nitrocellulose membranes (Optitran, AM0014-1) using wet transfer in 1X transfer buffer (25 mM Tris base, 192 mM glycine, 20% methanol) at 90 V for 3 h at 4°C with constant stirring and cooling. Following transfer, membranes were equilibrated in TBST (50 mM Tris-HCl pH 7.6, 150 mM NaCl with 0.1% Tween-20) and blocked in 5% (w/v) nonfat dry milk (Research Products International 61948-92848) in TBST for 1 h at room temperature. Membranes were washed three times (5 min each) with TBST and incubated with primary antibodies diluted in 5% BSA (Sigma-Aldrich, A9647-100G) in TBST for overnight (12/14 hrs) in the cold room. After primary incubation, membranes were washed three times with TBST and incubated with appropriate secondary antibodies diluted in blocking buffer (5% BSA in TBST) for 1 h at room temperature. Membranes were washed again three times with TBST and imaged using the LI-COR Odyssey imaging.

#### Septin assembly and sedimentation assays

Septin complex assembly into higher order multimers was performed by first dialyzing purified SEPT2/6/7/9 complexes overnight in BRB80 buffer (80 mM PIPES-KOH pH6.9, 2 mM MgCl2, 1mM EGTA pH 6.9) supplemented with 300 mM KCl. Subsequently, complexes were diluted to a concentration of 400 nM in BRB80 with either 300 mM KCl (high salt control) or 45 mM KCl (low salt) and 1 mM GTP, and were incubated at 4°C for 2 hours. Reactions were centrifuged at 18,500 RPM (39,000xg) with a fixed-angle TL100.3 rotor for 15 min at 25°C in a Beckman Coulter Optima TLX ultracentrifuge. Pellets were resuspended in the same buffer and volume with the corresponding supernatants. Equal volumes were analyzed by 10% SDS PAGE gel, which were stained with Coomassie Brilliant Blue. Gels were imaged on the Odyssey scanning system (LICOR) and quantified using densitometric analysis.

#### In vitro TIRF microtubule- and actin-binding assays Microtubule-binding assays

Taxol-stabilized microtubules were prepared by incubating unlabeled (80%), HiLyte647 (10%) and biotin conjugated (10%) porcine brain tubulin (Cytoskeleton Inc) in BRB80 (80 mM PIPES pH 6.9, 1 mM EGTA pH 6.9, 2 mM MgCl2, 10% (v/v) glycerol) supplemented with 1 mM GTP at 37°C for 35 minutes. The concentrated microtubule mix was incubated at 37°C for 20 minutes following the addition of 10 μM Taxol. Following polymerization, microtubules were pelleted by ultracentrifugation using a Beckman Coulter Optima TLX ultracentrifuge equipped with a TLA-100.3 fixed-angle rotor. Samples were centrifuged at 75,000 rpm for 30 min at 25°C. After centrifugation, the supernatant containing unpolymerized tubulin was carefully removed, and the microtubule-containing pellet was gently resuspended in 14 µL of microtubule stabilization buffer consisting of 80 mM PIPES pH 6.9, 1 mM EGTA pH 6.9, 2 mM MgCl₂, 10% (v/v) glycerol, supplemented with 1 mM GTP, 0.2 mM DTT, and 10 µM taxol to maintain microtubule stability. Resuspended microtubules were then kept at room temperature while protected from light.

Acid washed glass coverslips (0.15 mm thick) were mounted on a glass slide using double sided tape to create 8-10 μL motility chambers. To adhere biotinylated microtubules to the glass each chamber was incubated with 5 mg/mL biotinylated BSA (Sigma) followed by 0.5 mg/mL Neutravidin (Thermo Fisher) which was diluted in cold PBS. Chambers were incubated with taxol-stabilized microtubules diluted in BRB80 containing 1mM GTP for 15 min, followed by blocking buffer (BRB80, 1 mg/mL BSA, 1% w/v Pluronic F-127 (Sigma), 10 μM taxol) for an additional 5 min. Chambers were then washed with 80-100 μL HEPES buffer (30 mM HEPES-KOH pH 7.4, 50 mM of KOAc pH 7.4, 2 mM of MgOAc, 1 mM of EGTA-KOH pH 7.4 and 10% glycerol). GFP-SEPT9_i1, GFP-SEPT2/6/7/9 (WT, S82A/S85A, S82E/S85E) were diluted in the HEPES based septin dilution buffer containing HEPES, 0.1% w/v Pluronic F-127, 0.1 mg/mL BSA and 10 μM Taxol before applying to the chambers and incubated for 15 mins at room temperature. A final wash with 80-100 μL HEPES buffer was done before imaging.

#### In vitro actin binding assays

Filamentous actin (F-actin) was polymerized in vitro from unlabeled monomeric rabbit skeletal muscle actin (G-actin; Cytoskeleton Inc., AKL99). G-actin (28 µM) was incubated in F-actin polymerization buffer (Cytoskeleton Inc., BSA02-001) containing 10 mM Tris-HCl (pH 7.5), 2 mM MgCl₂, 50 mM KCl, 1 mM ATP, 5 mM Guanidine Carbonate (pH 7.5), and 4 mM DTT for 20 min at room temperature to induce filament formation. Polymerized F-actin was subsequently stabilized and fluorescently labeled by supplementation with iFluor647-phalloidin (Abcam, ab176759) and incubated for an additional 30 min at room temperature. Labeled F-actin was then kept at room temperature while protected from light.

Flow chambers (8–10 µL) were assembled by mounting acid-washed glass coverslips (0.15 mm thickness) onto glass slides using double-sided tape. Imaging chambers were incubated with 1% (w/v) Pluronic F-127 for 5 min to prevent nonspecific surface adsorption and subsequently washed with BRB80 buffer (80 mM PIPES, pH 6.9, 1 mM EGTA, pH 6.9, 2 mM MgCl₂). Chambers were incubated with 5 mg/mL biotinylated bovine serum albumin (biotin-BSA; Sigma-Aldrich, A8549) for 5 min, washed with BRB80, and then incubated with 0.5 mg/mL NeutrAvidin (Invitrogen, A2666) for an additional 5 min. Chambers were subsequently blocked with BRB80 containing 1% Pluronic F-127 and 1 mg/mL BSA for 5 min. Polymerized filamentous actin was diluted to 0.25 µM in BRB80 and introduced into the flow chambers, where it was allowed to bind to the surface for 20 min. Recombinant GFP-SEPT9_i1 and GFP-SEPT2/6/7/9 complexes (wild-type, phosphonull S82A/S85A, or phosphomimetic S82E/S85E) were clarified prior to use by ultracentrifugation at 50,000 rpm for 10 min at 4°C using a Beckman Coulter Optima TLX ultracentrifuge equipped with a TLA-100.3 rotor to remove aggregated or degraded protein. The supernatant was transferred to a fresh tube and diluted in BRB80 to the desired concentration.

Flow chambers containing immobilized F-actin were washed with BRB80 supplemented with 1 mg/mL BSA and incubated with recombinant GFP-septin proteins for 20 min. Chambers were then washed with 30 µL BRB80 prior to imaging.

#### In vitro actin bundling assays

Filamentous actin (F-actin) was polymerized from 28 µM monomeric human platelet G-actin (Cytoskeleton Inc., #APHL99) in polymerization buffer (10 mM Tris-HCl pH 7.5, 2 mM MgCl₂, 50 mM KCl, 1 mM ATP, 5 mM Guanidine Carbonate, and 4 mM DTT) for 20 min at room temperature. Polymerized F-actin was subsequently stabilized and fluorescently labeled by supplementation with iFluor647-phalloidin (Abcam, ab176759) and incubated for an additional 30 min at room temperature. The F-actin was then diluted to 2.8 µM in F-buffer (20 mM HEPES pH 7.4, 1 mM MgCl₂, 0.5 mM ATP, 4 mM DTT). To initiate binding, 1 µL of 2.8 µM F-actin was incubated with 500 nM GFP-SEPT9 (WT, S82A/S85A. S82E/S85E) for 15 min at room temperature in a reaction mixture containing 0.1% Pluronic, 0.1% Casein, and 1 mg/mL BSA. Flow chambers (8-10 µL) were assembled using acid-washed coverslips and incubated with 1% Pluronic F-127 for 5 min, followed by a BRB80 wash (80 mM PIPES pH 6.9, 1 mM EGTA, 2 mM MgCl₂). Surfaces were functionalized by sequential 5-min incubations with 5 mg/mL biotin-BSA (Sigma-Aldrich, #A8549) and 0.5 mg/mL NeutrAvidin (Invitrogen, #A2666), then blocked with 1% Pluronic F-127 and 1 mg/mL BSA in BRB80. The actin-septin mixture was introduced into the chamber, incubated for 15 min, and visualized via TIRF microscopy.

#### Microscopy and image processing

##### Super-resolution (SoRa) spinning disk confocal microscopy

U2OS and primary hippocampal neurons (Figures 1D, 2A-B, 2H, 6F, S1A, S1C-D) were imaged with a Nikon TiE2 motorized SoRa super-resolution spinning disk confocal microscope (Nikon Instruments, Tokyo, Japan) equipped with a Yokogawa CSU-W1 SoRa optical unit and a Hamamatsu ORCA-Fusion BT back-thinned camera. A 60×/1.42 NA oil immersion objective lens (Nikon Apo Plan) was used for all imaging unless otherwise noted. Excitation was achieved with 405, 488, 547, and 647 nm laser lines, and emission was collected sequentially using appropriate band-pass filters and dichroic mirrors to minimize spectral crosstalk. Imaging parameters (laser power, exposure time, detector gain, pinhole/conjugate field settings, and sampling frequency) were held constant within an experiment and across experimental conditions to enable quantitative comparison of fluorescence intensities. Z-stacks were acquired at 0.2 µm axial steps at a voxel size of ∼108 nm x 108 nm x 200 nm (xyx) to capture the full cellular volume. Stacks were denoised and deconvolved in NIS-Elements (Richardson-Lucy, 20 iterations) prior to quantification.

##### Wide-filed deconvolution microscopy

Imaging of fixed MDCK cells was performed with a NIKON Ti2E motorized widefield/deconvolution microscope system equipped with a Hamamatsu ORCA-Fusion BT back-thinned camera, Plan Apochromat Lambda D 60x oil immersion objective lens (N.A. 1.42) with phase contrast and DIC condensers and prisms, and a SPECTRA III illumination source for excitation at 390 nm, 440 nm, 475 nm, 510 nm, 555 nm, 575 nm, 637 nm and 748 nm wavelengths and band pass filters. Images were acquired with the Nikon Elements software with post-processing (denoising; LIM 2D/3D deconvolution) and analysis tool on a HP Z4 advanced imaging workstation. Samples were illuminated under conditions optimized to prevent photobleaching while ensuring adequate signal-to-noise ratio. Image stacks were acquired with 0.1–0.2 µm z-step intervals to capture the full three-dimensional structure of the specimen. Images were denoised and deconvolved, as indicated, using the Nikon Elements software.

##### TIRF microscopy

TIRF imaging of in vitro assays was performed on a Nikon Eclipse Ti2 inverted microscope equipped with an iLAS laser illumination system for TIRF, a CFI TIRF 60X oil-immersion objective (NA 1.49), and a high-sensitivity Kineti <22 Teledyne Photometrics-based camera for fluorescence detection, controlled with Nikon NIS-Elements software. Images were acquired with 100 ms exposure times and laser power at ∼10%.

#### Image Analysis and Quantifications

##### Quantification of septin colocalization with actin filaments and microtubules

Quantification of endogenous SEPT9 colocalization with actin filaments in single confocal optical sections (Figures 1E, 2C-F) was performed as follows. Briefly, the cell of interest was first cropped from the original image, and the channels were separated using Color > Split Channels. Background signal was subtracted with Process > Subtract Background, and filaments were segmented based on fluorescence intensity using Image > Adjust > Threshold using an empirically determined threshold range, which was applied uniformly to all images independently of experimental condition. Each mask was registered as an individual ROI, and a new ROI that corresponds to the colocalizing/overlapping regions between SEPT9 and actin filaments was derived using the AND function of Fiji. Using the Analyze Particles command of Fiji, the surface areas of the actin mask and the actin-Sept9 overlap mask were automatically derived, and the overlap area was calculated as a percentage of its corresponding actin filament area.

Quantification of SEPT9 colocalization with microtubules in 3D volume stacks of optical slices (Figures 1G and 2G) was performed with a custom Python pipeline (sept9_mt_coloc_3d.py) using Python 3.14.4 on Windows 11, with numpy 2.4.4, scipy 1.17.11, scikit-image 0.26.02, pandas 3.0.2, nd2 0.11.3, tifffile 2026.4.11, and matplotlib 3.10.83. The pipeline contains an optional CuPy GPU path for Gaussian filtering, morphology, and a custom analytic-eigenvalue 3-D Frangi filter; for the present study all steps were executed on CPU. All anisotropic 3-D operations were scaled by the measured z/xy voxel ratio so that physical scales (µm) were preserved. A cell mask was computed by triangle thresholding a Gaussian-smoothed sum of the two channels, followed by 3-D ellipsoidal closing and rejection of small components. A nuclear mask was derived inside each cell from a “diffuse-SEPTIN9, low-MT” score, nuc_score = smooth(SEP) × (1 − smooth(MT)), triangle-thresholded; candidate components were retained only if median MT signal inside the component was below a fixed fraction of the cell-wide MT median, preventing weak-MT cytoplasm from being misclassified as nucleus. All headline metrics were computed on the cell-minus-nucleus volume (exNuc). For each field the slope α of SEPTIN9 on tubulin was estimated as the median of (SEP − bg) / (MT − bg) over the lowest-quartile SEPTIN9 voxels within exNuc (those least likely to carry true SEPTIN9 signal). α was capped to a physically plausible bleed range and rejected above an upper bound, the latter regime indicating diffuse cytoplasmic SEPTIN9 rather than spectral bleed. Where applied, SEP_corr = max(0, SEP − α·MT). SEPTIN9 was background-subtracted with a 3-D ellipsoidal white top-hat. Microtubules were high-pass filtered by subtracting a Gaussian-blurred copy, suppressing centrosomal flare while preserving filament cores. A 3-D Frangi vesselness4 was computed at three scales using analytic Cardano eigendecomposition of the Hessian. Filament masks were obtained by hysteresis thresholding of the Frangi response within the cell mask, closed, restricted to exNuc, and 3-D skeletonised by the Lee algorithm5. Skeleton components below a minimum physical length were discarded. Inside exNuc we computed two metrics: (i; Figure 1H) The 3-D skeleton-overlap fraction — the fraction of SEPTIN9 skeleton voxels that lie within a 2-voxel dilation of the MT mask. The dilation (≈ one lateral PSF FWHM, ∼220 nm at 108 nm/px) provides a colocalization tolerance buffer accounting for diffraction-limited sub-pixel offsets between the SEPTIN9 and microtubule channels; (ii; Figure 2G) Manders coefficient — the fraction of total SEPTIN9 intensity lying within the microtubule mask, calculated as Σ I SEP9(MT mask) ⁄ Σ I SEP9(cell − nucleus).

##### Quantification of astral microtubule number and length, and septin-containing fraction

Astral microtubule length and number were quantified from immunofluorescence images using ImageJ (Fiji) by manually tracing individual astral microtubules with the segmented line tool. The number of microtubules per each centrosome was recorded, and the length of each microtubule was measured in Fiji, and averaged for each aster per cell. The fraction of microtubules that associated with septins was derived by scoring individual astral microtubule for presence of GFP-SEPT9_i1, SEPT2 and SEPT7 on at least half of their total length. The number of microtubules that contained septins was divided by the total number of microtubules per aster.

##### Quantification of microtubule- and actin-bound septins

Quantification of fluorescence was performed in Fiji/ ImageJ using the plot profile function. A plot line with a width of 1 was drawn in individual microtubules or actin having a length of at least 10 µm. A straight line with same width was plotted as the background and was about 10 µm in length and 2 µm away from any microtubules or actin. The mean fluorescence intensity of each microtubule/actin was plotted after subtracting the mean fluorescence intensity of the background.

##### Quantification of number and length of actin bundles

The 488 nm SEPT9 channel was extracted from the raw two-channel image. A multiscale Sato vesselness filter (sigmas = 1.5, 2.5, 4, 6, 8 px; bright ridges) was applied to enhance tubular structures and suppress isotropic puncta and aggregates. The ridge-response image was threshholded at median + 5 × 1.4826 × MAD (≈ 5 σ above the noise floor; MAD = median absolute deviation). MAD-based thresholding was chosen over Otsu because Otsu was unreliable on images containing very bright outlier puncta. Foreground objects smaller than 15 px were removed. Single actin filaments were determined to have a mean width of ∼2–3 pixels, and actin bundles were defined as filaments with a width of ≥ 12 pixels, representing a 3-fold increase over single filaments due to crosslinking of multiple single filaments. Segments shorter than the average lenth of single filaments (10 µm) were discarded. Bundle-class components additionally had to satisfy aspect ratio ≥ 3 to be retained, in order to reject round aggregates. The number and length of each actin bundle was measured in image fields of equal surface area.

##### Quantification Sept9 colocalization with tubulin, F-actin and pSer9-GSK3β in neurons

Image segmentation was performed in a custom Python pipeline. A neurite mask was generated from a thresholded composite of the F-actin and Sept9 channels (Otsu threshold, scaled by 0.35; small components < 100 px removed). Growth cones were identified as terminal bulb-like expansions at neurite tips (area > 5 µm², circularity < 0.7) and dilated by 3 px to recover labile membrane edges. The axon shaft was defined as the elongated, high-aspect-ratio neurite region proximal to each growth cone. Every FOV contributed one growth-cone-averaged and one shaft-averaged measurement per readout. Pearson’s correlation coefficient was computed pixel-wise on the background-corrected intensities within each compartment (whole growth cone and axon shaft) for every colocalization partner pair: SEPTIN9 vs acetylated α-tubulin, SEPTIN9 vs F-actin, and SEPTIN9 vs pSer9-GSK3β. One Pearson r per FOV, per compartment, per partner pair was carried forward to statistics.

All inferential statistics were performed in GraphPad Prism 11 (v11.0.0.84; GraphPad Software, San Diego, CA, USA). For each colocalization partner and each compartment, vehicle vs CHIR-99021 per-FOV Pearson values were compared by the two-tailed Mann–Whitney U test (α = 0.05). Six comparisons were performed in total — SEPTIN9–acetylated-tubulin, SEPTIN9–F-actin and SEPTIN9–pSer9-GSK3β, each evaluated separately in growth cone and in axon shaft — and each pairwise test is reported individually. Cohen’s d was computed alongside as a standardized effect size [(CHIR mean − control mean) / pooled SD] and reported descriptively (small ≈ 0.2, medium ≈ 0.5, large ≈ 0.8).

##### Quantification of neurite asymmetry

Neuronal polarity was assessed using the Single Neurite Tracer (SNT) plugin in Fiji. Neurite lengths were traced, and polarized neurons were classified as neurons with a single neurite, which was at least twice in length as the second longest neurite.

##### Statistical analyses

Statistical analysis of data and graph generation was performed with GraphPad Prism 11 statistical analysis software (v11.0.0.84; GraphPad Software, San Diego, CA, USA). Scatter plots display the individual values of all data sets, and indicate the mean values plus/minus standard error of the mean (SEM). Data set groups were first checked for standard deviation (SD) variance using the Prism’s Descriptive Statistics tool and then, assessed for normal distribution of variance using the D’Agostino and Pearson and Shapiro-Wilk normality tests. For pair-wise comparisons of data with normal distributions, a student’s t test was used if SDs were equal, or a Welch’s t-test if SDs were unequal. The Mann-Whitney U-test was used for non-normally distributed data. P-values were derived and statistical significance was set at *p* < 0.05 as the standard threshold against the null hypothesis; *p* values > 0.05 were deemed as not significant (ns).

## Supplementary Figures

**Figure S1.**
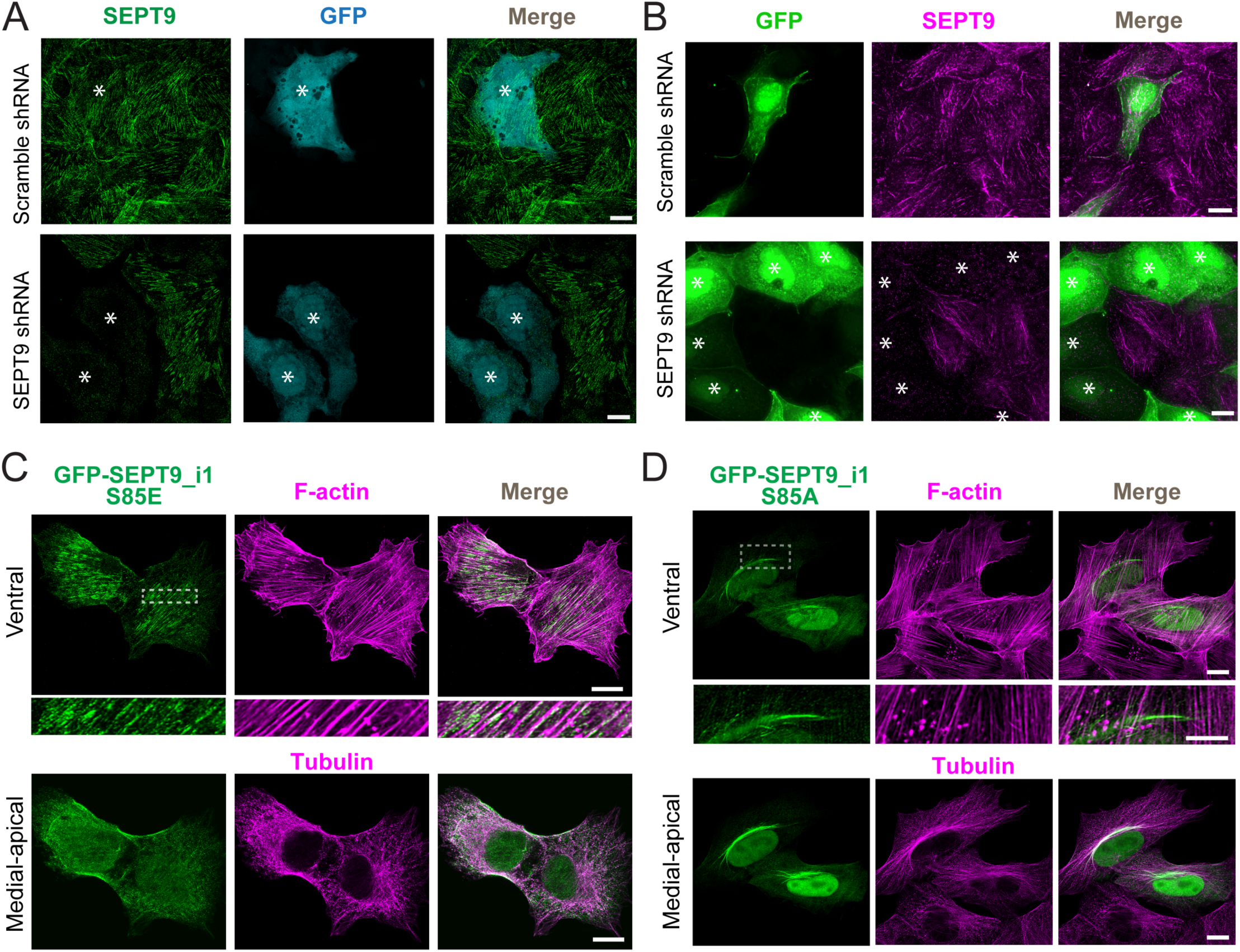
Endogenous SEPT9 depletion, and localization of the SEPT9_i1 mutants S82E and S82A in MDCK cells. **(A)** Super-resolution SoRa spinning-disk confocal microscopy images show maximum intensity projections of ventral optical sections of U2OS cells transfected with plasmids expressing GFP (cyan) and scramble control (top) or SEPT9 (bottom) shRNAs, and stained for endogenous SEPT9 (green). Asterisks indicate cells transfected with GFP-expressing plasmids. Scale bars, 10 μm. **(B)** Denoised and deconvolved wide-field fluorescence microscopy images show maximum intensity projections of medial-apical optical sections of MDCK cells, which were transfected with plasmids expressing GFP (green) and scramble control or SEPT9 shRNAs, and stained for endogenous SEPT9 (magenta). Asterisks indicate cells transfected with GFP-expressing plasmids. Scale bars, 10 μm. **(C-D)** Super-resolution SoRa spinning-disk confocal microscopy images show maximum intensity projections of ventral (top) and medial-apical (bottom) optical sections of U2OS cells transfected with GFP-SEPT9_i1-S85E (green; C) or GFP-SEPT9_i1-S85A (green; D), stained for actin (phalloidin; top in magenta) and α-tubulin (bottom in magenta). Dashed rectangles indicate regions shown at higher magnification. Scale bars, 10 μm.

**Figure S2.**
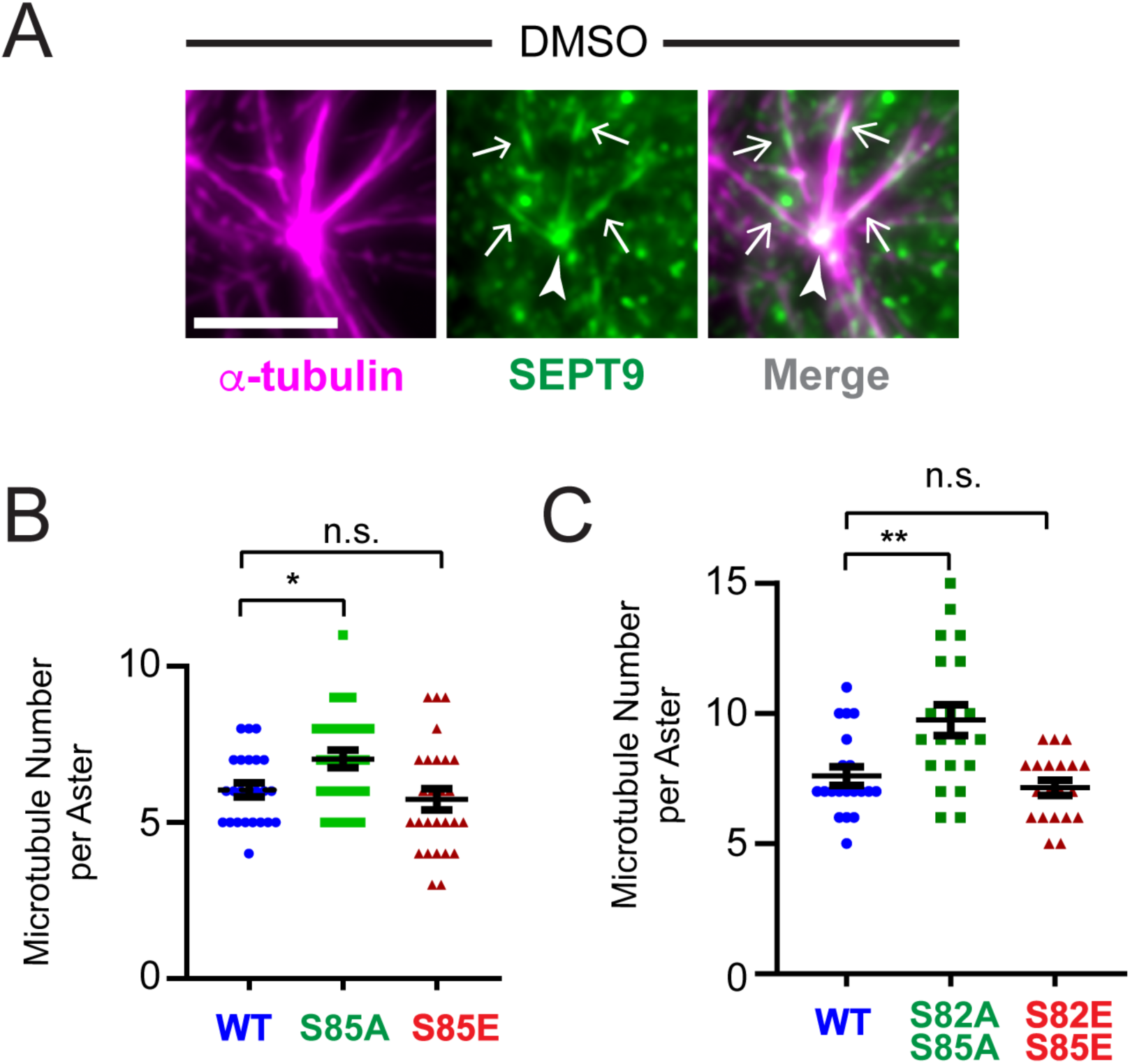
Endogenous SEPT9 localizes to centrosomal microtubules, and phosphonull mutations of SEPT9 at S82 and S85 increase the number of centrosomal microtubules. **(A)** Denoised and deconvolved wide-field fluorescence microscopy images show maximum intensity projections of medial-apical optical sections of MDCK cells, which were treated with nocodazole (1.6 μM) for 2 h at 37 °C, followed by 40 seconds nocodazole wash-out at 25 °C. Images show endogenous SEPT9 (green) and α-tubulin (magenta). Arrows point to SEPT9 on the lattice of astral microtubules and arrowhead points to SEPT9 at the centrosome. Scale bar, 10 μm. **(B)** Quantification of the number (mean ± SEM) of astral microtubules per aster following nocodazole wash-out in MDCK cells transfected with plasmids expressing SEPT9 shRNA and shRNA-resistant wild type GFP-SEPT9_i1, GFP-SEPT9_i1-S85A and GFP-SEPT9_i1-S85E. Pairwise comparisons between wild-type (*n* = 24 cells), S85A (*n* = 28 cells), and S85E (*n* = 27 cells) mutants were statistically analyzed using a Mann-Whitney U-test. *, p < 0.01; n.s., not significant. **(C)** Quantification of the number (mean ± SEM) of astral microtubules per aster following nocodazole wash-out in MDCK cells (*n* = 20) transfected with plasmids expressing SEPT9 shRNA and shRNA-resistant wild type GFP-SEPT9_i1, GFP-SEPT9_i1-S82A/S85A and GFP-SEPT9_i1-S82E/S85E. Pairwise comparisons between wild-type, S82A/S85A, and S82E/S85E mutants were statistically analyzed using a Mann-Whitney U-test. **, p < 0.01; n.s., not significant.

**Figure S3.**
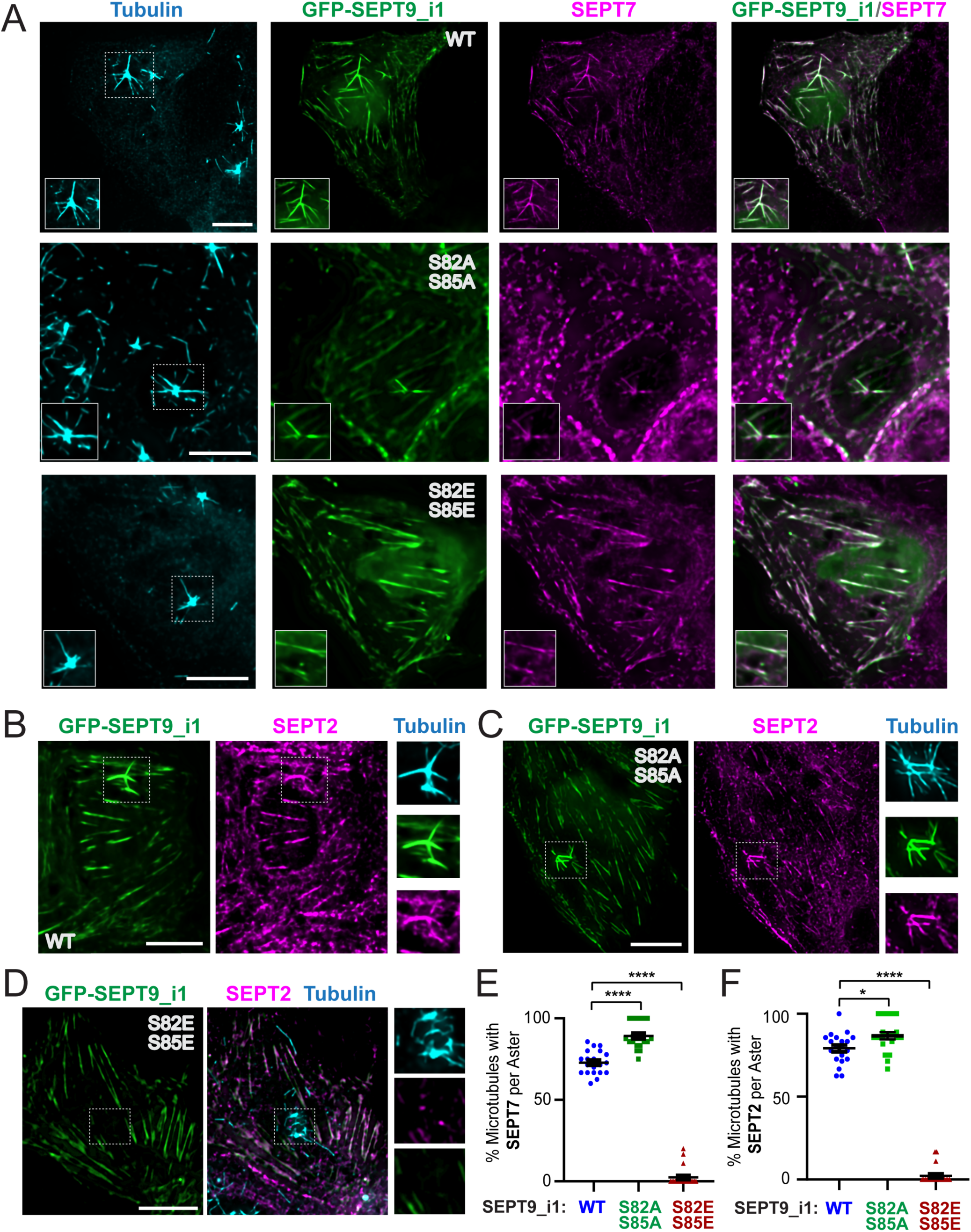
Phosphonull mutations of SEPT9_i1 at S82 and S85 enhance localization of endogenous SEPT7 and SEPT2 to centrosomal microtubules, whereas phosphomimetic mutations have the opposite effect. **(A)** Denoised and deconvolved wide-field fluorescence microscopy images show maximum intensity projections of medial-apical optical sections of MDCK cells, which were transfected with plasmids expressing SEPT9 shRNA and shRNA-resistant GFP-SEPT9_i1 (wild-type and indicated phosphomutants; green), and stained for endogenous α-tubulin (cyan) and SEPT7 (magenta). Cells were treated for nocodazole (1.6 μM) for 2 h at 37 °C followed by washout for 40 seconds at 25 °C to allow centrosomal microtubule regrowth prior to fixation and staining. Insets show magnified views of regions outlined by dashed rectangles. Scale bars, 10 μm. **(B-D)** Denoised and deconvolved wide-field fluorescence microscopy images show maximum intensity projections of medial-apical optical sections of MDCK cells, which were transfected with plasmids expressing SEPT9 shRNA and shRNA-resistant wild type GFP-SEPT9_i1 (B), GFP-SEPT9_i1-S82A/S85A (C) and GFP-SEPT9_i1-S82E/S85E (D), stained for endogenous α-tubulin (blue) and SEPT2 (magenta). Cells were treated for nocodazole (1.6 μM) for 2 h at 37 °C followed by washout for 40 seconds at 25 °C to allow centrosomal microtubule regrowth prior to fixation and staining. Insets show magnified views of regions outlined by dashed rectangles. Scale bars, 10 μm. **(E)** Quantification of the percentage (mean ± SEM) of astral microtubules decorated with endogenous SEPT7 per aster following nocodazole wash-out. Pairwise comparisons between wild-type (*n* = 20 cells) and S82A/S85A (*n* = 20 cells) or S82E/S85E (*n* = 20 cells) mutant were statistically analyzed using a Mann-Whitney U-test. ****, p < 0.0001. **(F)** Quantification of the percentage (mean ± SEM) of astral microtubules decorated with endogenous SEPT2 per aster following nocodazole wash-out. Pairwise comparisons between wild-type (*n* = 20 cells) and S82A/S85A (*n* = 20 cells) or S82E/S85E (*n* = 20 cells) mutant were statistically analyzed using a Mann-Whitney U-test. *, p < 0.05; ****, p < 0.0001.

**Figure S4.**
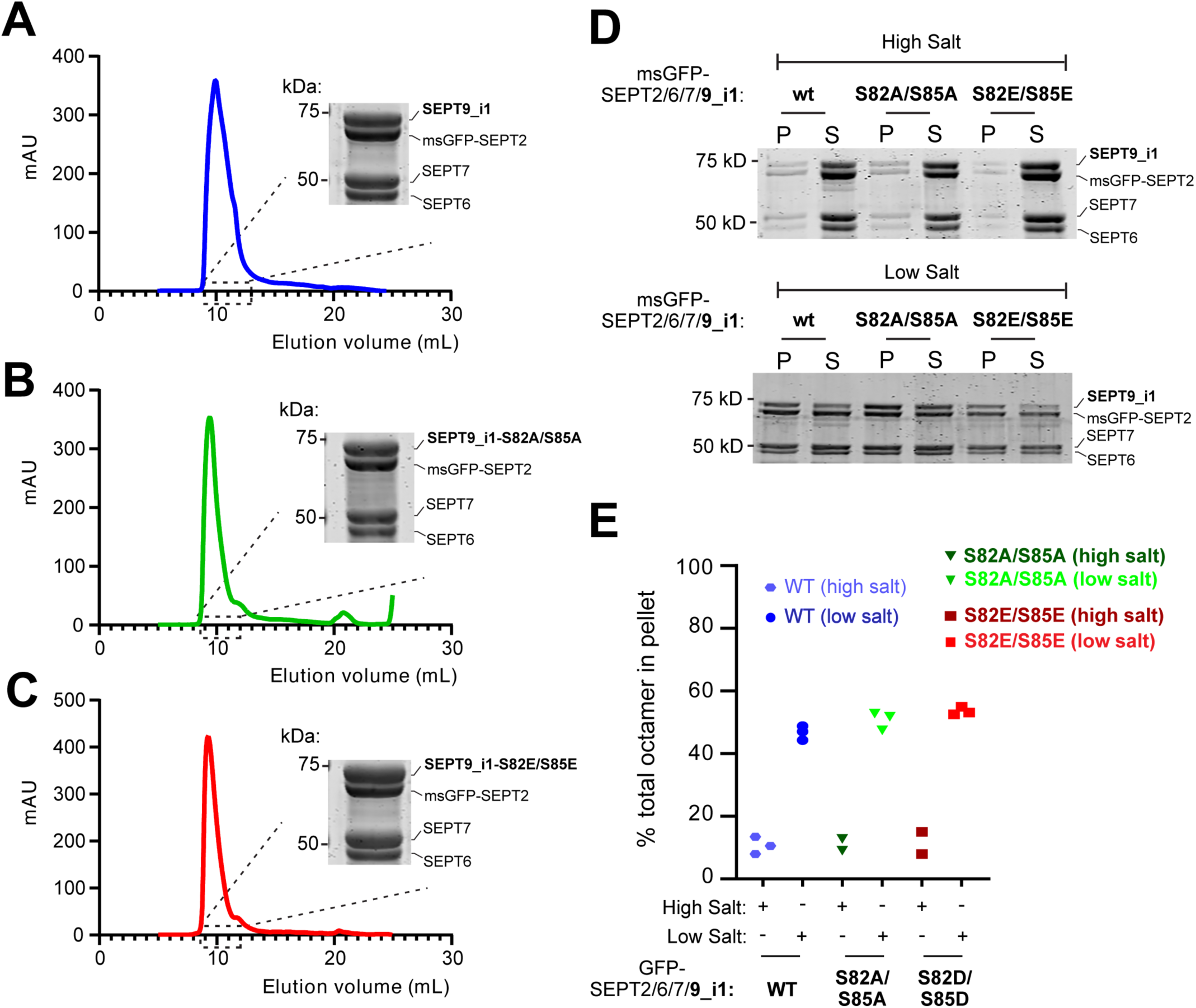
Phosphonull and phosphomimetic mutation of SEPT9_i1 at S82/S85 do not impair assembly into SEPT2/6/7/9 oligomers and polymers. **(A-C)** Size-exclusion chromatography elution profiles — absorbance at 280 nm (mAU) as a function of elution volume — of msGFP-tagged SEPT2/6/7/9_i1 complexes following dual tag affinity purification from bacterial lysates that contained Strep-tagged SEPT9_i1 (A), SEPT9_i1-S82A/S85A (B) or SEPT9_i1-S82E/S85E (C) along with SEPT6, SEPT7 and His-/msGFP-tagged SEPT2. Complexes were resolved on a Superdex 200 Increase 10/300 GL column. Coomassie Blue-stained SDS-PAGE gels show the protein composition of fractions pooled from the main elution peaks (dashed rectangles). **(D)** Coomassie Blue-stained SDS-PAGE gels of pellet and supernatant fractions from sedimentation assays of SEPT2/6/7/9_i1 complexes containing wild-type, S82A/S85A, or S82E/S85E SEPT9_i1. Complexes were diluted and incubated for 2 h in either high-salt buffer (300 mM KCl; polymerization control) or low-salt buffer (45 mM KCl), which induces assembly into higher-order polymers. Following sedimentation at 39,000 × g, pellets were resuspended in a volume equal to that of the supernatant, and equal volumes of each fraction were resolved by SDS-PAGE. **(E)** Quantification of the fraction of total SEPT2/6/7/9_i1 complex protein recovered in the pellet following incubation in high- and low-salt buffer. Data are from three independent experiments in low-salt buffer and two to three independent experiments in high-salt buffer (control).

**Figure S5.**
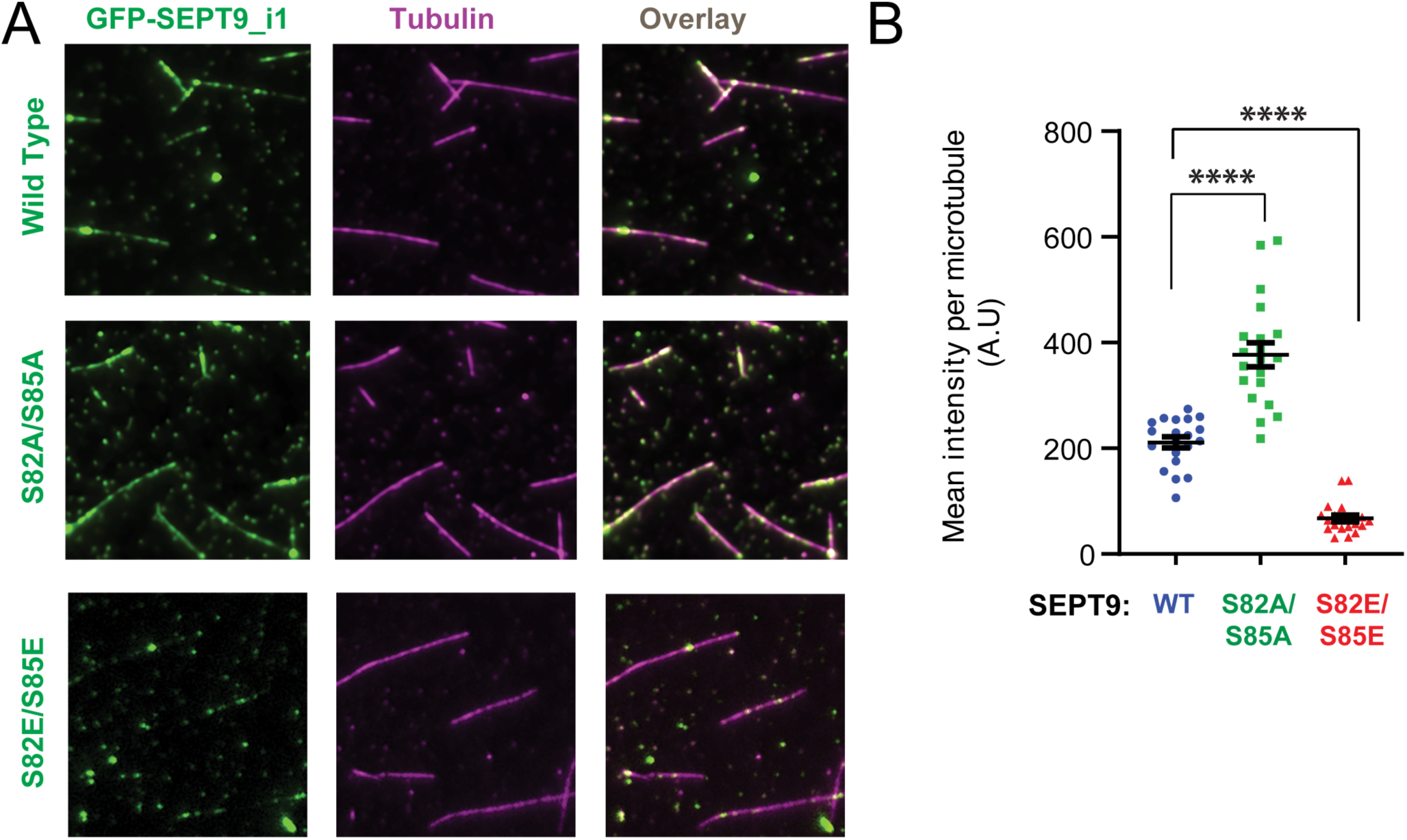
Phosphonull and phosphomimetic mutations of SEPT9_i1 at S82/S85 increase and reduce, respectively, binding to microtubules in vitro. **(A)** Total internal reflection fluorescence (TIRF) microscopy images of taxol-stabilized microtubules (magenta) following incubation with 3 nM GFP-SEPT9_i1 (green) for wild type, and S82A/S85A, and S82E/S85E mutants. Scale bars, 10 μm. **(B)** Quantification of mean GFP fluorescence intensity per microtubule length following incubation with 3 nM GFP-SEPT9_i1 (*n* = 20; WT, S82A/S85A, S82E/S85E). Pairwise comparisons between WT and S82A/S85A were statistically analyzed using an unpaired Welch’s t-test, and between WT and S82E/S85E with a Mann-Whitney U-test. ****, p < 0.0001 Plot shows mean ± SEM.

**Figure S6.**
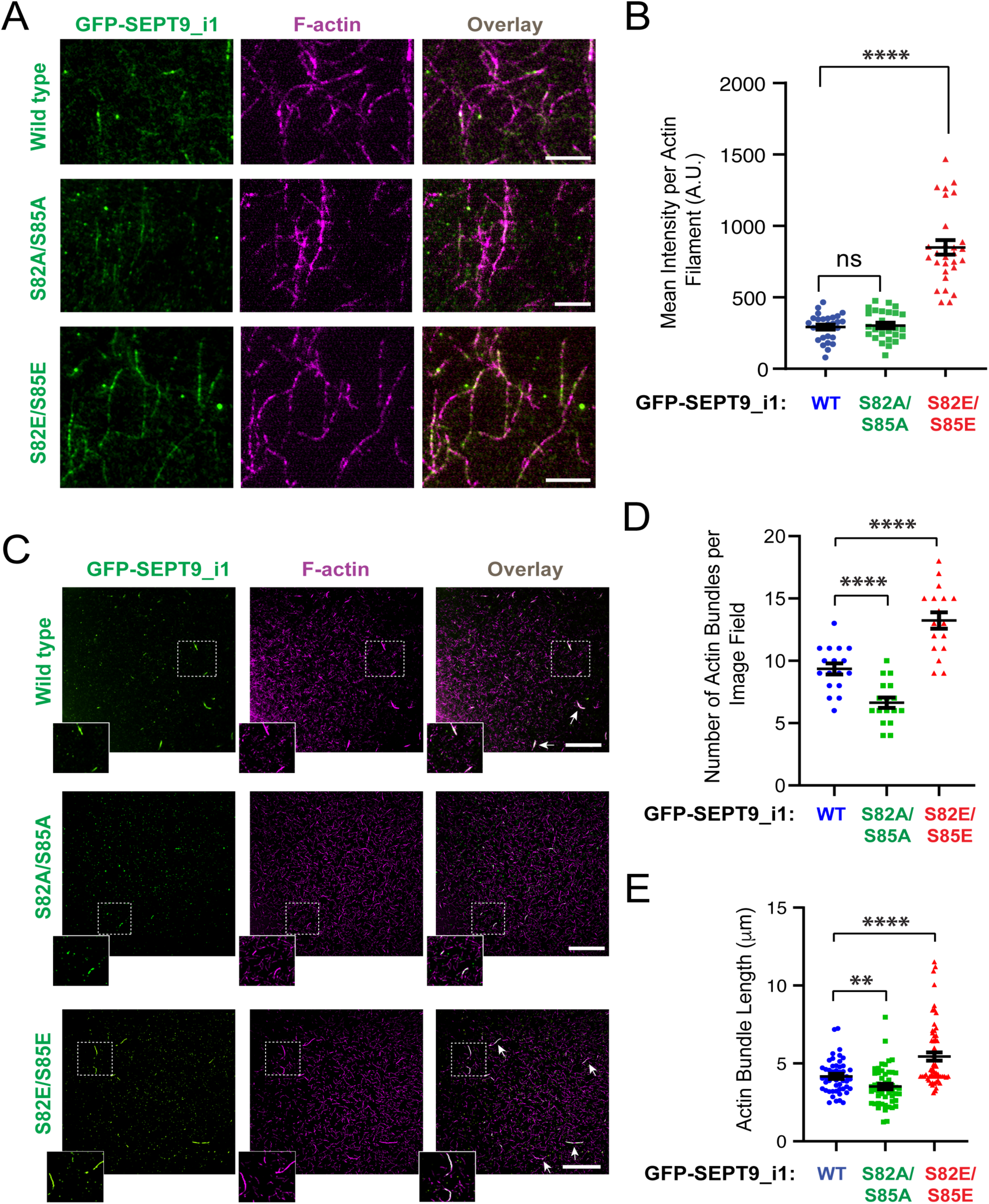
Phosphomimetic mutations of SEPT9_i1 at S82/S85 enhance actin binding and bundling in vitro, and phosphonull mutations decrease actin bundling. **(A)** Total internal reflection fluorescence (TIRF) microscopy images of phalloidin-stabilized actin filaments (magenta) following incubation with 200 nM GFP-SEPT9_i1 (green) for wild type, and the S82A/S85A and S82E/S85E mutants. Scale bars, 5 μm. **(B)** Quantification of mean GFP fluorescence intensity per actin length following incubation with 200 nM GFP-SEPT9_i1 (*n* = 28; WT, S82A/S85A, S82E/S85E). Pairwise comparisons between WT and S82A/S85A was statistically analyzed with a Welch’s t-test, and between WT and S82E/S85E with a Mann Whitney U-test. ns, not significant; ****, p < 0.0001 **(C)** Total internal reflection fluorescence (TIRF) microscopy images of phalloidin-stabilized actin filaments (magenta), which were flown into the TIRF chamber following an incubation with 500 nM GFP-SEPT9_i1 (green; wild type, S82A/S85A or S82E/S85E) in solution for 15 minutes at 25 °C. Insets show at higher magnification regions with actin bundles outlined with dashed rectangles. Arrows point actin bundles. Scale bars, 20 μm. **(D)** Quantification of the number (mean ± SEM) of actin bundles per image field (*n* = 17) of equal surface area for GFP-SEPT9_i1, GFP-SEPT9_i1-S82A/S82A and GFP-SEPT9_i1-S82E/S85E. Pairwise comparisons between WT and the mutants were statistically analyzed using an unpaired Welch’s t-test. ****, p < 0.0001 **(E)** Quantification of the length (mean ± SEM) of actin bundles for GFP-SEPT9_i1 (*n* = 48), GFP-SEPT9_i1-S82A/S82A (*n* =50) and GFP-SEPT9_i1-S82E/S85E (*n* = 61). Pairwise comparisons between WT and the mutants were statistically analyzed using the Mann-Whitney U-test. **, p < 0.01; ****, p < 0.0001

